# Rab11 negatively regulates Wingless preventing JNK mediated apoptosis in *Drosophila* epithelium during embryonic dorsal closure

**DOI:** 10.1101/2021.04.27.441306

**Authors:** Nabarun Nandy, Jagat Kumar Roy

## Abstract

Cell signaling pathways involved in epithelial wound healing, show a lot of complexities when it comes to their regulation. Remarkably, a large proportion of these signaling pathways are triggered at the time of morphogenetic events which usually involve epithelial sheet fusions during embryonic development, such as the event of dorsal cloure in Drosophila embryos. One such conserved pathway in the wound healing process is the JNK-Dpp signaling pathway. Recent observations suggest that one such upstream regulator of JNK mediated apoptosis could be Rab11, a small Ras like GTPase, which is functionally associated with the membrane and cortical cytoskeletal organization of epithelial cells. Using *Drosophila* embryonic dorsal closure as a model of wound healing, we observed that a targeted expression of a *Rab11* loss of function mutant in the dorso-lateral epidermis of fly embryos (tissue which extends contra-laterally in order to fill the intervening gap) undergoing dorsal closure leads to an ectopic expression of Caspase-3 and a concomitant up-regulation of the JNK-Dpp signaling. This resulted in the death of the dorso-lateral epithelial cells with a consequent embryonic lethality due to dorsal closure defects. Interestingly, a simultaneous knockdown of *wingless* (another developmentally conserved gene) in *Rab11* mutants resulted in a rescue of the lethal phenotype and also a significant level of successful completion of the dorsal closure process. In our experiments we suggest Rab11 could promote cross talk between the JNK-Dpp pathway and the canonical *wingless* pathway in the regulation of apoptosis in the dorsolateral epithelium of fly embryos undergoing dorsal closure.

**One Sentence Summary:** Rab11 functions through a conserved Wingless mediated JNK-Dpp pathway during embryonic dorsal closure.

## Introduction

Dorsal closure (DC) and epithelial morphogenesis in *Drosophila* embryos provides deep insights into the properties of epithelial cells required for the genesis of a completely differentiated epithelial tissue, and also, its maintenance upon a damage (Wood et al, 2002 and Kiehart et al, 2017). Cellular properties involved in this process includes coordinated cell shape changes which eventually establishes a continuum of epithelial cells on the dorsal side of the embryo, which, in the initial stages of dorsal closure remains covered by a single layer of polyploid epithelial cells known as the amnioserosa (Lawrence and Morel, 2003). The feature shows striking resemblances with the process of epithelial wound healing. The properties of cells involved in tissue repair include sensing of the changes in mechanical properties of the cell membrane (Vogel and Sheetz, 2006; Ilina and Freidl, 2009) upon a damage which is then transduced in the form of intra-cellular and intercellular signaling (Heisenberg and Bellaiche, 2013). The cellular processes involved in *Drosophila* embryonic dorsal closure share several parallels with wound healing in many a physiological and cell-biological aspects (Martin and Parkhurst, 2004) where changes in individual properties of epithelial cells have an impact on the morphologies of entire group/cohort of cells involved in the dorsal closure process. Unique among these properties are cell-cell adhesion dynamics, oscillations of cortical cytoskeletal elements and most importantly spatio-temporal expression of morphogens which help cells communicate as they show coordinated shape changes while closing the dorsal opening. Here we were interested to dissect the regulation and roles of signaling pathways in the cells of the dorso-lateral epithelium (DLE) of DC undergoing embryos emphasizing upon the regulation of the conserved JNK-Dpp mediated apoptotic pathway which has also been rigorously reported to be regulating the physiology of cells surrounding an epithelial wound enabling them to fill it up (Bosch et al, 2005).

The exact effects of the activation of JNK pathway seems to be unclear, however, JNK’s regulatory effects on both pro and anti-apoptotic cascades (Jing and Anning, 2005; Kolahgar et al, 2011), in developmental morphogenesis (Agnes et al, 1999; Stronach and Perrimon, 2002), tumor progression and metastasis (Zhu et al, 2010) have been reviewed over the years. The JNK pathway has been reported to be regulating the process of collective cell migration, wound healing and associated inflammation (Friedl and Gilmour, 2009; Ramet et al, 2002). Genes involved in JNK signaling which regulate the events of *Drosophila* “Dorsal Closure” and “Thorax Closure” are critically important as reports of Jacinto et al, 2002 and Martin-Blanco et al, 2000 suggest. Further, reports of Hou et al, 1997 and Fernandez et al, 2007, state that JNK expression in the dorsal most epithelial cells (DMEs) is essential for the release of Dpp morphogen in the DMEs bringing about coordinated cell shape changes by triggering downstream signaling events (Widmann and Dahmann, 2009).

Thus organization of membrane associated molecules which essentially regulate downstream cell signaling events such as the JNK pathway is of a prime necessity to epithelial cells, which they achieve through the endomembrane system. Here the small ras like GTPases, Rabs, play an instrumental role as they mark distinct protein sorting, endomembrane compartments within the eukaryotic cell (Zerial and Stenmark, 1993). These components traffic membrane associated proteins such as receptors or cell-cell adhesions which function as important cellular domain markers and therefore decide cell polarity, and also stabilize the cortical cytoskeleton of epithelial cells which is essential for the maintenance of their cognate morphologies and also cell signaling events. Thus Rab11, a marker of the Recycling Endosomes, which has been reported to be an essential regulator of the spatio-temporal oscillations of cell-cell adhesion, cytoskeletal and cell surface receptors dynamics in epithelial cells likely plays critical roles in dorsal closure as well as epithelial wound healing process.

*Rab11* plays an instrumental role in the development and differentiation process of several epithelial organs of *Drosophila* such as the embryonic epithelium (Sasikumar and Roy, 2009), the eye (Alone et al, 2005) and even gonads (Tiwari and Roy, 2008). Tiwari and Roy, 2009 and Bhuin and Roy, 2010 have shown that Rab11 seemed to be regulating the eye and the wing developmental processes *via* a common JNK signaling pathway. The association of Rab11 functions with the JNK pathway in eye or wing morphogenesis intrigued us to look into Rab11’s probable roles on the regulation of the JNK-Dpp signaling pathway in the dorso-lateral epithelia (DLEs) of *Drosophila* embryos during dorsal closure as DLEs show a persistent JNK-Dpp signaling during the closure event.

Here, through a targeted perturbation of Rab11 functions, we show that *Rab11* plays an instrumental role in the physiology of the dorsolateral epidermis of fly embryos undergoing DC. Our experiments involve a detailed analysis of the effects of the targeted expression of the *Rab11* LOF mutant, i.e., *Rab11^N124I^* in the physiology of the DLEs of embryos undergoing DC with a probable mechanism of action.

## Results

### Functional loss of Rab11 in the dorsolateral epidermis results in the failure of dorsal closure of *Drosophila* embryos

In order to study the effects of disruption of Rab11 function during DC, a *UAS-Rab11^N124I^*, i.e., *Rab11^DN^* (Roeth et al, 2009) stock was driven with *pnr-Gal4* in the DLEs (Kushnir et al, 2017) and, to ensure the specific expression domain of *pnr-Gal4*, a *UAS-GFP* transgenic stock was driven with it, where, green fluorescence specific to the dorso-lateral epidermis (Calleja et al, 2000) of embryos was observed (Fig1. A-B’). Thus, the expression domain of *pnr-Gal4* was considered ideal to study the effects of targeted *Rab11* perturbation. The embryos wherein *Rab11* had been knocked out in the DLEs showed a conspicuous dorsal opening shown in the white box of Fig 1-E-E’ suggesting that the conditional expression of a *Rab11* LOF mutation in the DLEs results in a failure of dorsal closure. Cuticle preparations of *pnr-Gal4* driven *UAS-Rab11^DN^* embryos revealed that out of a total 150 mutant cuticles observed, all showed dorsal closure defects, anterio-posterior compression/shortening and ventral puckering suggesting the mutant phenotype is absolutely penetrant. Embryonic lethality assay results shown in Fig. 1F corroborates the cuticular defects seen in the mutants, where a mean embryonic lethality of 24.9±1.6% was recorded (supplementary table T1). The *pnr-Gal4* driven *UAS-Rab11^DN^* mutants also phenocopied the *puckered* (a *hep* phosphatase and a negative regulator of activated JNK (Martin Blanco et al, 1998)) homozygous mutants as shown in Fig.1 G-J. We further analyzed effects of *EP3017* (Rab11 LOF) homozygotes on the epithelial morphogenetic process. Lethality assay revealed an 8.6 ± 0.9% (Table T7) embryonic lethality of *EP3017* homozygous mutant embryos. Although *EP3017* homozygous mutants do not emerge as third instars and show a lethality of 17.9±0.7% lethality as first instar larvae, which interestingly also show dorsal anterior lesions (shown in white arrows in supplementary Fig. S1-A) in the thoracic segments along the dorsal midline. Out of the total dead *EP3017* homozygous embryos, 4.2 ± 0.3% of the embryos showed dorsal closure defects as shown in supplementary Fig. S1-B. These observations confirm that Rab11 functions in the DLEs are indispensable for dorsal closure in *Drosophila* embryos.

**Fig. 1.**
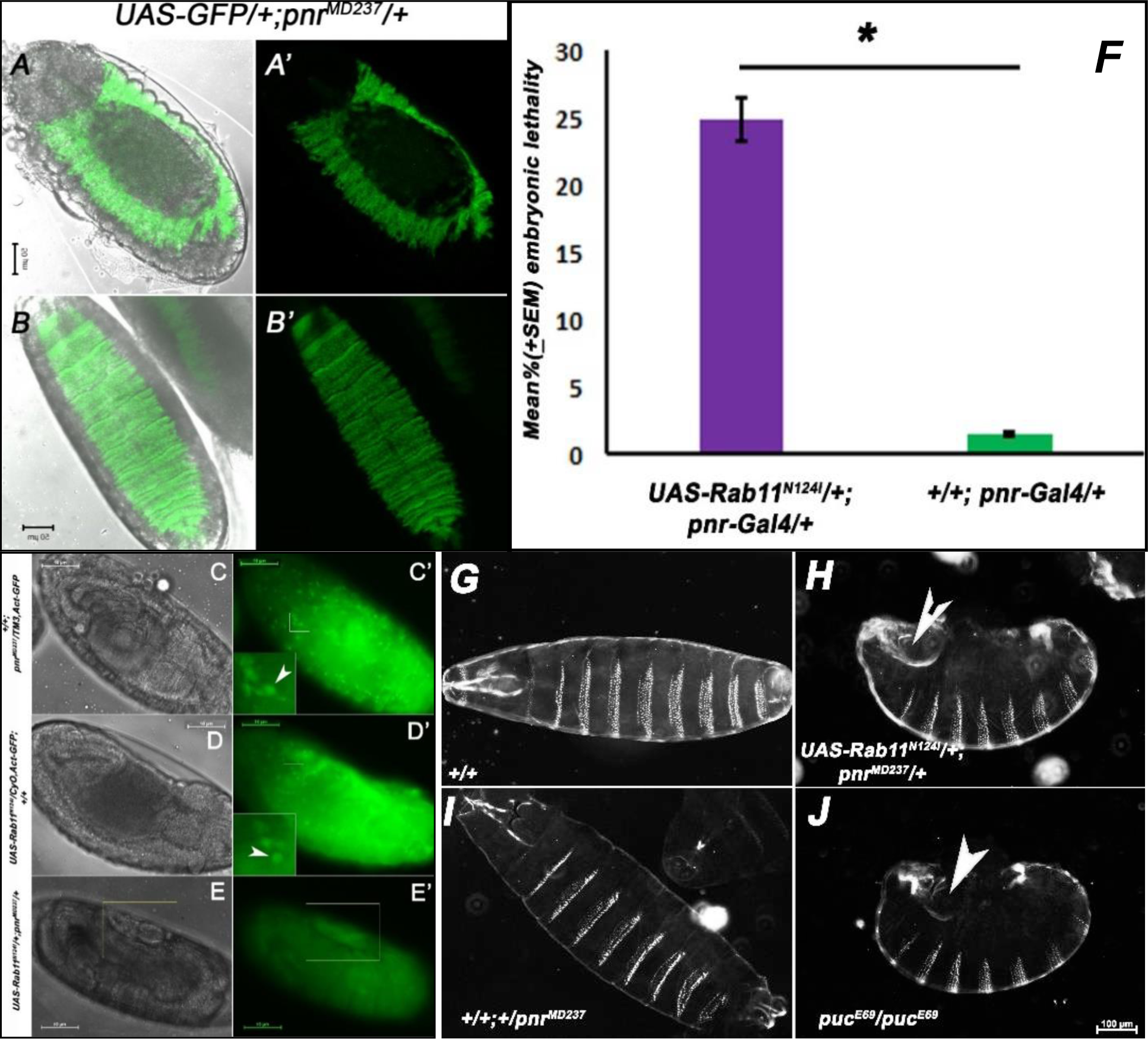
Conditional loss of Rab11 in the DLEs results in an incomplete dorsal closure. Phase contrast and fluorescence images of *pnr-Gal4*(*pnr^MD237^*) driven *UAS-GFP* embryos (A-B’)during and after dorsal closure stage showing the expression domain of *pannier* promoter.(C-E’) Phase contrast and fluorescence image of driver(C-C’, *pnr-Gal4/TM3-Act-GFP*), responder(D-D’, *UAS-Rab11^N124I^/CyO-Act-GFP*) and driven(E-E’, *UAS-Rab11^N124I^/+; pnr-Gal4/+)* embryos (Insets in C’ and D’ showing the expression of Actin-GFP in actin rich haemocytes (shown with white arrows) which helped detect the presence of balancer chromosomes in the driver and responder stocks and consequently the driven embryos in which the balancers have segregated out.(F) Graph representing Mean (± SEM) % embryonic lethality in a *pnr-Gal4* driven *UAS- Rab11^N124I^*genetic background as compared to undriven controls.(G-J) Dark field images of 22- 24h developed embryonic cuticles where *pnr-Gal4* driven *UAS-Rab11^N124I^*individuals show strong phenotypic resemblances with *puckered* homozygous mutants such as puckering of the cuticle, anterior-posterior shortening and a conspicuous dorsal opening, a stereotypical phenotype also shown in the white box of E-E’.

### Perturbation of Rab11 function in the DLEs results in their failure to attain proper morphology required for dorsal closure

The dorsal closure defects intrigued us to look into the effects of *Rab11* mutation on the individual morphologies of the DLEs as *Rab11* mutants fail to undergo DC. Therefore, stage 13- 14 embryos, were fixed and immunostained for the membrane marker FasIII and Rab11 (Fig 2- A-C’). Confocal sections of the undriven embryos, revealed that the DLEs and also the LEs, during DC, undergo a substantial amount of dorso-ventral elongations (Inset in Fig.2 A’). At the same time Rab11 too appeared as discrete punctae, in the cytoplasm of the LEs and DLEs (Fig. 2 B’). Confocal sections also revealed that Rab11 co-localizes considerably with FasIII (Shown in white arrows in the inset of C’) suggesting its probable role in the maintenance of basolateral junctional integrity of the epithelial cells which is lost in *pnr-Gal4* driven *UAS-Rab11^DN^* embryos (Fig. 2 –D-F’). Confocal images of embryos revealed that the *Rab11* mutant DLEs failed to attain the dorso-ventrally elongated morphology Fig. (2 D’). Also, Rab11 expression in the DLEs as well as the LEs of the mutant embryos was found to be diffused and scattered unlike the controls, suggesting a loss of the spatial expression pattern of Rab11 which is a necessary characteristic for its domain specific functions within a cell.

**Fig. 2.**
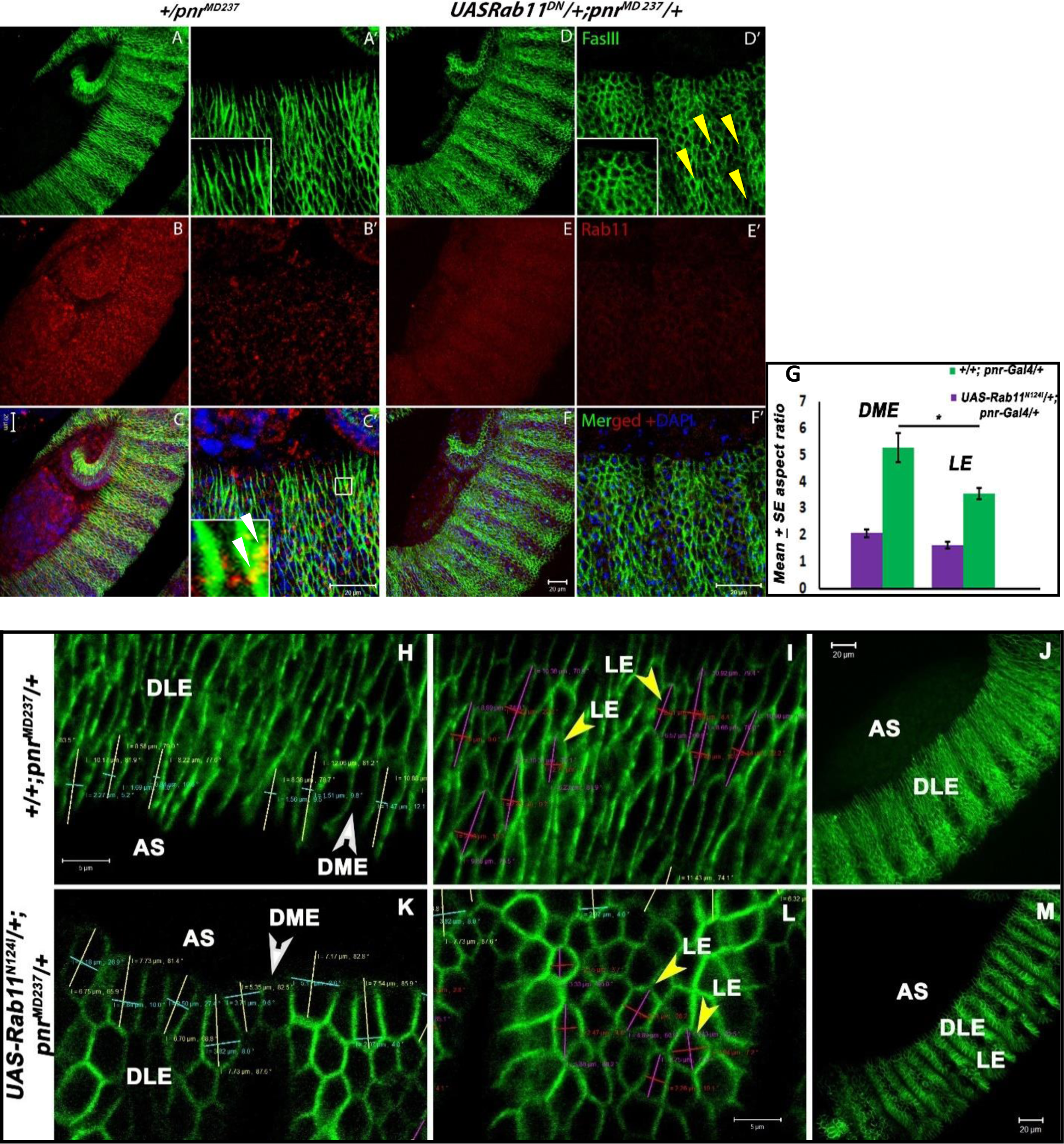
A Functional loss of Rab11 in the DLES results into an aberrant cell shape. (A-C) Confocal projection images of stage 13-14(dorsal closure) stage *pnr^MD237^/+,* undriven embryos immunostained for FasIII(Green), Rab11(Red) and counterstained with DAPI (Blue). (A’-C’) Enlarged confocal sections of the LE of embryo shown in A-C. Inset in A’ shows the dorso- ventrally elongated morphology of DMEs and DLEs which is a consequence of amnioserosal retraction and constriction of the actin purse string at the PAS/DME interface. Inset in C’ shows Rab11 significantly co-localizes with the baso-lateral tight junction marker FasIII, shown with white arrow. (D-F) Confocal projection images of stage 13-14 stage, *pnr-Gal4* driven *UAS- Rab11^N124I^* embryos immunostained for FasIII, Rab11 and DAPI. The distribution of Rab11 in the LE is diffused and random in E unlike the punctate expression of Rab11 in B. (D’-F’) Enlarged confocal sections of the embryo shown in D-F. Inset in D’ shows the FasIII expression pattern of the DLEs and DMEs which are polygonal unlike the cells of the wild type embryo but the cells of the LE juxtaposed to the DLEs towards the ventral side of the embryo show the usual rhomboid and dorsoventrally elongated morphology much like the wild type cells shown by yellow arrow. Panels from H-I and K-L show the enlarged confocal sections of the projection images of stage 13-14, un-driven and *pnr-Gal4* driven *UAS-Rab11^N124I^*, embryos shown in panels J-M. Yellow and blue lines in H-I represent the length and breadth of the DMEs and the same (length and breadth) are represented by magenta and red lines for the LEs shown in I-L. Graph in panel G represents the mean aspect ratio of the DMEs and LEs of the undriven controls (Green bars) with respect to the *pnr-Gal4* driven *UAS-Rab11^N124I^*individuals (Violet Bars). Paired T test was performed to assess the significance between the comparable classes and a P-value ≤0.05 was considered as significant (n=100).

Mean aspect ratio values of the DLEs were calculated as shown in Fig 2 H-I and K-L and graphs were plotted for both DMEs and LEs in the *pnr-Gal4* driven *UAS-Rab11^DN^* condition as well as in the controls where significant differences were observed (Fig. 2 G).

### Loss of *Rab11* function in the dorso-lateral epidermis of DC undergoing embryos triggers apoptosis in the DLEs and results in a membrane associated expression of Caspase-3

The loss of cellular elongations in *Rab11* mutant conditions intrigued us to observe the fate of the DLEs in *Rab11* mutants where, approximately in all (98±0.67%) embryos, a characteristic dorsal opening with a conspicuous necrotic patch on the periphery of the opening was evident (Fig 8. J-J’- shown in white arrow) indicating a probable apoptosis of these cells. Thus stage 13- 14 embryos were immunostained for Caspase-3 and FasIII, where, the expression of Caspase-3 represents a Caspase-9 like activity of Dronc in *Drosophila* (Fan and Bergmann, 2010). It was observed that in *Rab11* mutant DLEs, a significant increase of Caspase-3 expression followed as compared to the controls (Fig. 3 A-I and Fig. 3 A’-L’). We performed the same experiment on *EP3017* homozygous embryos where we found that a 2.9 ± 0.19 % (data not shown) of the homozygous mutant embryos showed a significant increase of Caspase 3 expression, although, the expression was more pronounced in the region of the DLEs, signifying Rab11 could be a negative regulator of apoptosis in *Drosophila* embryonic epithelium (Supplementary Fig S2 G-L). The elevated expression of Hid, in the DLEs of *pnr-Gal4* driven *UAS-Rab11^N124I^* individuals further confirmed our hypothesis (Supplementary Fig S2 A-F).

**Fig. 3.**
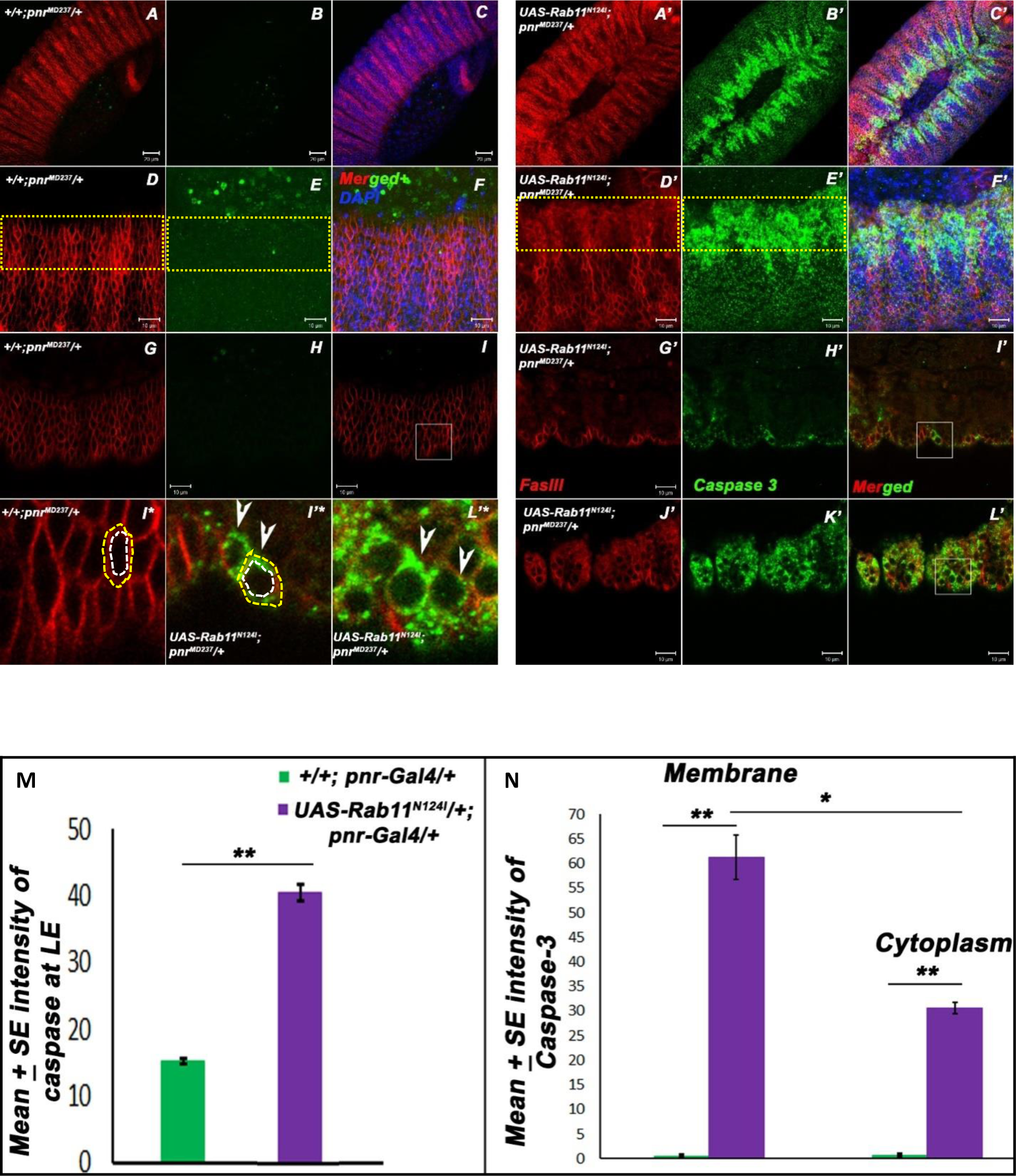
Perturbation of Rab11 function in the DLEs of DC undergoing embryos leads to an ectopic activation of Caspase-3. (A-C) Confocal projection images of a dorsal closure stage (stage 13-14) (*pnr^MD237^/+*) embryo immunostained for FasIII (Red), Caspase-3 (Green) and counterstained with DAPI (Blue). (D-F) 2.5 times magnified image of the lateral epidermis of the embryo shown in A-C, revealing the distribution of Caspase-3 which appears as speckles spread over the cytoplasm. Panels G-I shows a single section/slice of the lateral epidermis shown in D-F. (A’-C’) Confocal projection images of a dorsal closure stage (stage 13-14) of *pnr-Gal4* driven *UAS-Rab11^N124I^*embryo immunostained for FasIII (Red), Caspase-3 (Green) and counterstained with DAPI (Blue). (D’-F’) 2.5 times enlarged projection image of the lateral epidermis of the embryo shown in A’-C’, revealing the distribution of Caspase-3 which appears as speckles spread over the cytoplasm. G’-I’ and J’-L’ panels show individual sections of the lateral epithelium shown in D’-F. I*, I’* and L’* are 4 times magnified images of I, I’ and L’ showing the distribution of Caspase expression in the epithelial cells. The yellow and white dotted lines around the cells mark the outer periphery of the membrane and the outer periphery of the cytoplasm respectively, and the area enclosed between the two has been considered for the analysis of membrane associated Caspase expression. (M) Graph representing the mean intensity of Caspase-3 in the DLEs of the *pnr-Gal4* driven *UAS-Rab11^N124I^*condition as compared to the undriven controls. Intensity was measured within the region enclosed by the yellow box shown in panels E and E’ (yellow boxes D and D’ denote the region of DLEs) for 30 embryos. A P value ≤ 0.01 was considered to be highly significant represented by **. (N) Graph comparing the mean intensity of Caspase-3 on the plasma membrane of the mutant cells vs the undriven ones and also the cytoplasm of mutant cells versus the undriven ones. N=25 cells of each genotype, where a P value ≤0.05 was considered significant and a P value ≤0.01 was considered to be highly significant.

Analysis and quantification of Caspase-3 in *Rab11* mutant DLEs (shown in Fig 3I* and Fig 3 I’*) revealed that membrane and cytoplasmic expression of Caspase-3 was significantly more in the *pnr-Gal4* driven *UAS-Rab11^N124I^* embryos as compared to the un-driven controls (Graph N (Fig3)). It was also observed that the membrane associated Caspase-3 expression in *pnr-Gal4* driven *UAS-Rab11^DN^* condition was higher than the cytoplasmic expression levels in the same embryos suggesting significant role of Caspase-3 on the membrane, and a potential role could be the proteolytic cleavage of membrane associated components such as cell-cell adhesions, receptors or cytoskeletal anchorage proteins.

### Cortical actin and Myo-V organization is perturbed in the DLEs on a targeted disruption of Rab11 functions in flyembryos

The morphological abnormalities in the DLEs of Rab11 mutants further intrigued us to observe the organization of the cortical actin cytoskeleton/cortactin in these cells as actin based adhesions dynamics are critical to morphogenesis (Jacinto et al, 2000). Thus *Rab11* mutant embryos were immunostained for phosphorylated tyrosines present in the Actin nucleation centers (ANCs) of the cortical cytoskeleton (Fig. 4 H-J’). Interestingly it was found that ANCs localize to the cellular cortex of the control DLEs (Fig 4 H-H’), however, in *EP3017* homozygotes, a diffusion of the cortex associated phosphotyrosines was clearly visible. The severity of this phenotype was even more pronounced in *pnr-Gal4* driven *UAS-Rab11^N124I^*embryos, suggesting *Rab11*’s essential role in the maintenance of the integrity of cortical actin cytoskeleton of the epithelial cells. Further investigation into the intracellular distribution of the Myosin V, a non-conventional myosin, which associates with the cortical actin (Aguilar-Aragon et al, 2020, Lo Presti et al, 2012) revealed that MyoV expressed fairly in the cortical region of the DLEs in control embryos, which is clearly evident from the partial overlap of Myosin expression with FasIII expression towards the cytosolic side of the DLEs shown in A’ and A” (marked in white arrows). However, in *pnr-Gal4* driven *UAS-Rab11^N124I^* embryos Myosin V expression was found to be much diffused, haphazard and reduced unlike their un-driven counterparts. Cumulatively, our results indicate that *Rab11* LOF in the DLEs result in the complete disruption of their apical domain by strongly affecting the cortical acto-myosin complexes which are critically involved in the epithelial cell shape plasticity as well as membrane organization during DC.

**Fig. 4.**
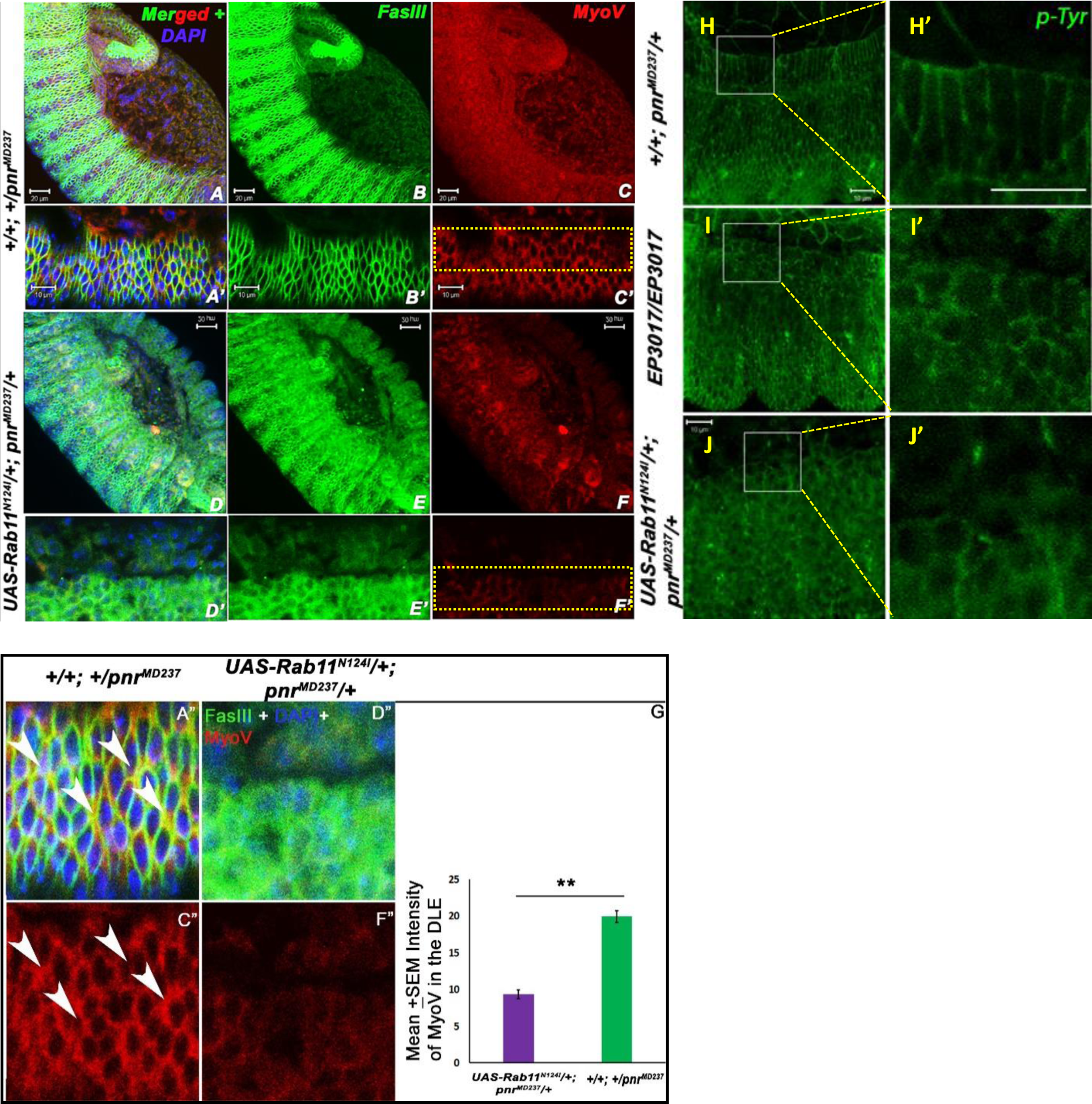
Cortical Acto-myosin complex integrity is lost in cells wherein Rab11 functions are perturbed. (A-C)Confocal projection images of*+/+; +/pnr^MD237^*, stage 13-14 embryos stained for FasIII(Green), Myosin V(Red) and counterstained with DAPI (blue). Panels A’-C’show 2.5 times magnified section of the LEs of the embryo shown in A-C. Panels D-F show confocal projections of *pnr-Gal4* driven *UAS-Rab11^N124I^*embryos immunostained for FasIII (Green), MyoV(Red) and DAPI(Blue) and corresponding sections are represented in panels D’-F’. Panels A” and C” show the 2.5 times magnified image of sections shown in panels A’ and C’. White arrows in A” show the regions of “near-overlap (seen as regions of yellow)” of MyoV with the inner boundary of FasIII justifying its cortical localization in the epithelial cells of the lateral epidermis which is disrupted in D” and F”. (G) Graph representing mean intensity of MyoV in the *pnr-Gal4* driven *UAS-Rab11^N124I^* condition vs undriven embryos where intensities were measured from 25 embryos from the region of the DLE as shown in C’ and F’. P value≤0.01 was considered to be significant. (H-J) Confocal projection images of the lateral epidermis of stage 13-14 embryos immunostained for phosphorylated Tyrosines (depicting the ANCs), where (H’-J’) represent their 4 times magnified sections. H-H’ represent the undriven condition where ANCs are localised on the cell boundary. The shape of these cells are obviously dorso-ventrally elongated, and the green fluorscence is restricted to the cell margin, justifying the cell shape. I-I’ represent the *EP3017/EP3017* mutant where ANCs are somewhat diffused suggesting a disruption of cortical actin cytoskeleton of the lateral epidermal cells and this diffusion is much more pronounced in *pnr-Gal4* driven *UAS-Rab11^N124I^* individuals(J-J’).

### A *Rab11* LOF in the DLEs of embryos undergoing DC leads to a cytoplasmic accumulation and elevated expression of Armadillo/β-Catenin

The cortical cytoskeleton organization maintains cell shape changes during epithelial morphogenesis and it is meticulously anchored at cell-cell adhesions such as the Adherens junctions. Roeth et al, 2009 have rigorously shown that DE-Cadherin is recycled via Rab11 to the membrane of the amnioserosa as well as lateral epidermis during DC. As Cadherins connect to the cytosolic cortical actin with the help of α and β-catenins, we were interested to look into the expression of the latter in the absence of Rab11 function, as β-catenin/Armadillo not only organizes the cortical actin at the adherens junction (White and Vincent, 1996), but also is an important component of the canonical Wingless signaling pathway. Thus, embryos undergoing DC were fixed and immunostained for Armadillo in an undriven, *pnr-Gal4* driven *UAS- Rab11^N124I^* and *EP3017* homozygous mutant embryos where in Fig. 5 A-C’, Armadillo in undriven embryos showed a rather membrane associated expression pattern along with a faint cytoplasmic expression too. In a *pnr-Gal4* driven *UAS-Rab11^DN^*condition it was observed that the expression of Armadillo was not only diffused in the DLEs but also elevated (Fig 5 B), at the same time, Armadillo was also found to be co-expressing with the nuclei of the DLEs (shown in the inset of Fig 5 B’ with white arrows). A similar elevation of Armadillo expression was also observed in the DLEs of 3.27±0.3% (data not shown) of the *EP3017* homozygous, DC defective embryos with a concomitant loss of membrane associated Armadillo expression throughout the embryonic epidermis.

**Fig. 5.**
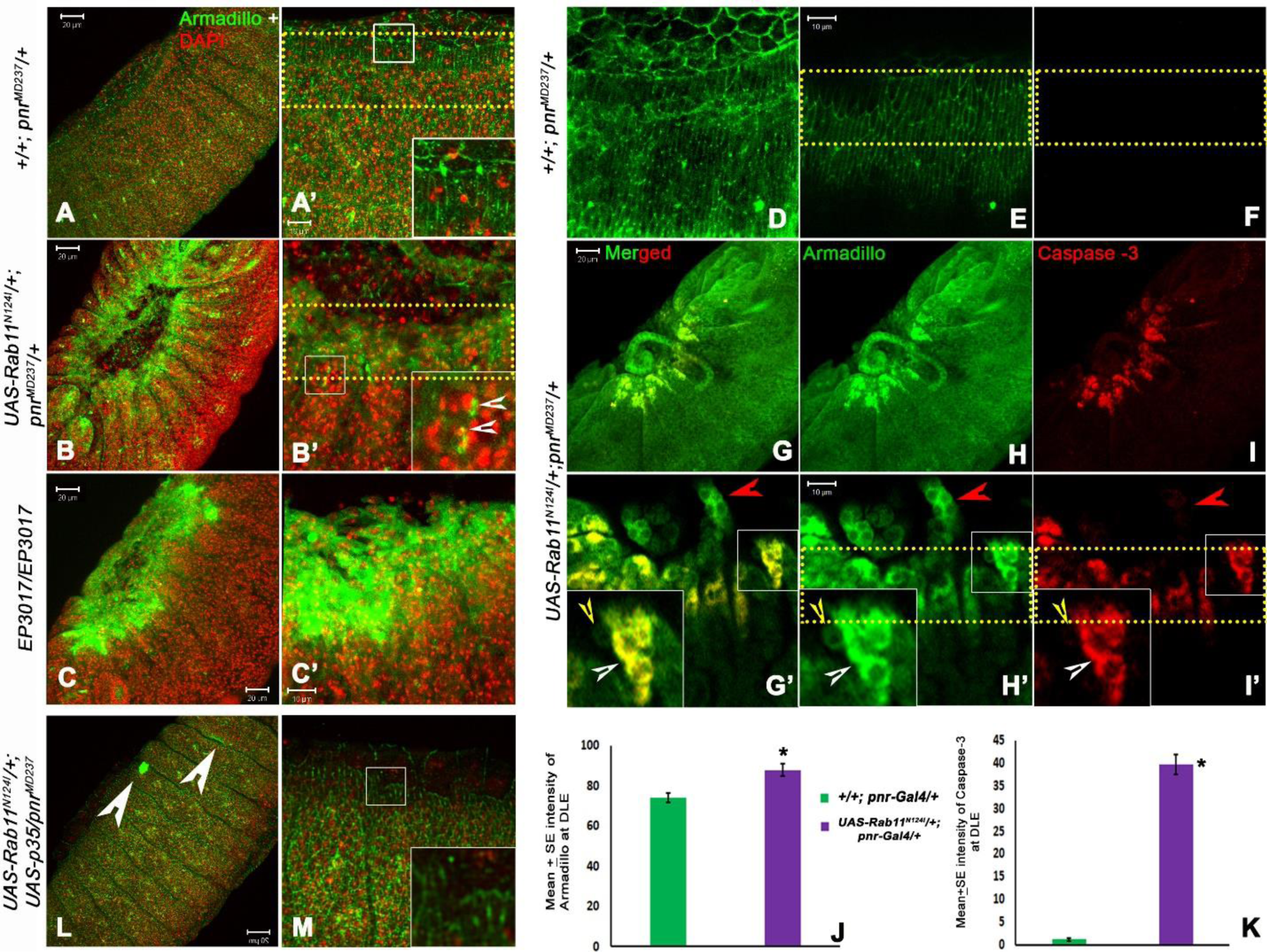
Disruption of Rab11 function in the epithelial cells results in cytoplasmic accumulation of β-Catenin/Armadillo. (A-C’) Confocal projection images of stage 13-14 developed embryos immunostained for Armadillo (Green), and counterstained with DAPI (Red), where A’-C’ show 2.5 times magnified sections of the images shown in A-C respectively. The mutant embryos shown in B-B’ and C-C’ show elevated Armadillo expression and a disruption of its membrane associated distribution as compared to A-A’. Inset in B’ represents the distribution of Armadillo in *pnr-Gal4* driven *UAS-Rab11^N124I^* condition where Armadillo co- localizes with the nucleus as shown by the white arrows. D shows projection image of undriven, stage 13-14 embryos immunostained for Armadillo (Green) and Caspase-3 (Red). E-F represent sections/slices of the same showing the distribution of Armadillo and Caspase 3 respectively. (G-I) are confocal projection images of *pnr-Gal4* driven *UAS-Rab11^N124I^*embryos which are again co-stained for Armadillo (Green) and Caspase-3 (Red) (Caspase 3 has been shown with Red which is a pseudo color in place of Far Red, and the choice of emission spectra of secondary antibodies was made with the purpose to avoid bleed through at the time of signal acquisition). G’-I’ show the 2.5 times enlarged section of the embryo shown in panels G-I. White arrows in the insets show the 2 times enlarged image of the DLE cells with elevated levels of Cytoplasmic Armadillo (Green) which completely co-localizes(Yellow) with elevated cytoplasmic Caspase3(Red). Yellow arrow shows an adjacent cell which does not have elevated Armadillo as well as lacks elevated Caspase-3 expression. Red arrow shows the membrane associated distribution of Armadillo in the embryonic Malpighian tubule cells where there is negligible expression of Caspase-3. (J) Graph showing the mean intensity of Green fluorescence in the regions shown within the dotted box in A’ and B’ and (K) Graph showing mean intensity of Caspase -3 in the area enclosed by the dotted boxes shown in panels F and I’. Intensities were measured for 25 embryos of each genotype and a P value ≤0.01 obtained from paired t-test between the comparable classes was considered significant. L and M represents the stage 13-14 *pnr-Gal4* driven *UAS-Rab11^N124I^; UAS-p35* embryo immunostained for Armadillo, where membrane its associated expression like the controls is somewhat restored suggesting a feeble rescue. White arrows in L represent regions of ectopic and elevated expressions of Armadillo in the *pnr-Gal4* driven *UAS-Rab11^N124I^; UAS-p35* embryos suggesting a partial rescue of the cytoplasmic and elevated Armadillo expression observed in *pnr-Gal4* driven *UAS-Rab11^N124I^* embryos.

As shown earlier, an elevation of Caspase-3 expression and its membrane associated distribution is a significantly stereotyped phenomenon in *Rab11* mutant embryos (Fig3). Interestingly, the pattern of Caspase expression was found to be much similar to Armadillo expression in the DLEs of the same mutant embryos, so, it was interesting to find whether or not Armadillo and Caspase-3 co-express in the same intra-cellular domains of the DLEs, which we presume to be indicative of a plausible interaction between the two. Thus *Rab11* mutant embryos were co-immunostained for Armadillo and Caspase-3 (Fig 5 D-I’) which revealed that Caspase-3 showed an overlap with Armadillo in the cytoplasm of the DLEs of Rab11 mutant embryos. We also found that, DLEs with elevated Caspase 3 expression are the ones which show an elevated expression of Armadillo on their membranes as well as cytoplasm (shown with white arrows in the insets of G’, H’ and I’), whereas, the other cells do not. Interestingly, the induction of a simultaneous expression of the anti-apoptotic gene p35, results in the feeble restoration of Armadillo expression (Fig 5 L-M) with, a not much significant rescue of embryonic lethality as observed in *pnr-Gal4* driven *UAS-Rab11^N124I^* embryos, signifying that Rab11 may not be solely responsible for Armadillo organization at the adherens junctions, although its roles in DE-Cadherin recycling is fairly established. The elevated Armadillo expression in *Rab11* mutants could be an outcome of an elevated Caspase activity where Caspase-3 specifically co-localizes with Armadillo, probably its proteolytic activity on the latter could be a reason of the dissociation of Armadillo from the Cadherin complex followed by its cytoplasmic accumulation thereby mimicking a “Wingless overexpressed” physiology.

### Conditional expression of *Rab11-*LOF mutation leads to the an ectopic EMT induction in the DLES with an up-regulation of MMP1 and Rho1 expression levels

The loss of spatio-temporal organization of cell-cell adhesions with a subsequent loss of apical actin-myosin arrangement could plausibly drive the *Rab11* mutant DLEs towards a mesenchymal fate from an epithelial one. MMP1 being a significant mesenchymal marker therefore intrigued us to observe the expression of the same in, stage 13-14 *Rab11* mutant embryos where we observed an elevated expression of MMP1 in approximately all {98.74 ± 0.8% (data not shown)} the *pnr-Gal4* driven *UAS-Rab11^DN^* embryos and also in 4.03±0.34% (data not shown) of the *EP3017* homozygous dead embryos (Fig 6 Graph P) corroborating our assumption.

**Fig. 6.**
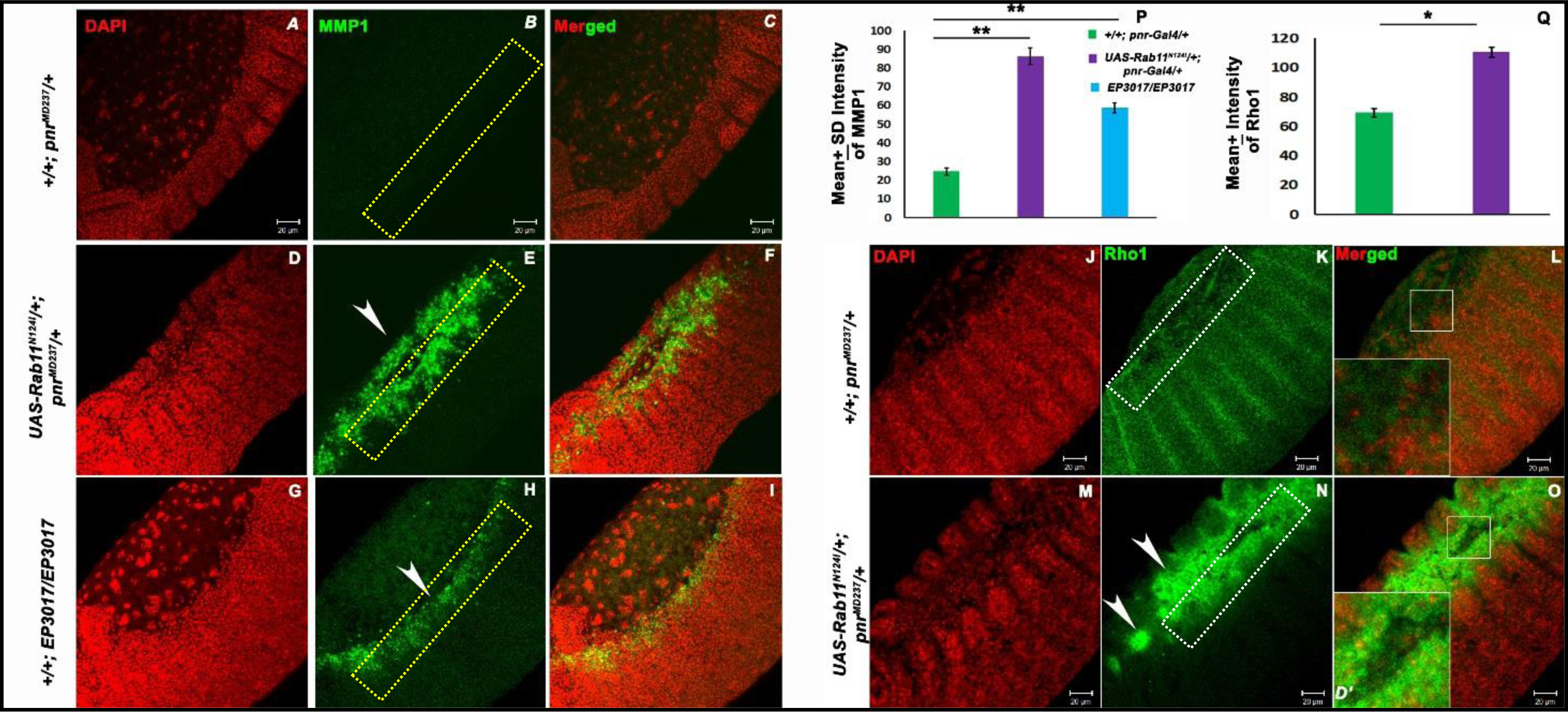
Cells wherein Rab11 functions are perturbed in the DLEs results in an ectopic induction of EMT along with elevated Rho1 signaling. (A-I)Confocal projection images of stage 13-14 embryos immunostained for MMP1(Green) and counterstained with DAPI(Red), where panels D-F represents a *pnr-Gal4* driven *UAS-Rab11^N124I^*embryo and panels G-I represents *EP3017* homozygous mutant embryos. MMP1 (Green) expression levels were measured in the area enclosed within the dotted square shown in the panels B-H for 25 embryos of each genotype and plotted in the graph P. White arrows in the afore-mentioned panels shows the regions of elevated MMP expression in the conditionally driven as well as *EP3017* homozygotes. (J-O) Confocal projection images of embryos undergoing dorsal closure immunostained for Rho1 in *pnr-Gal4* driven *UAS-Rab11^N124I^*embryos (M-O) as compared to undriven embryos (J-L). Insets in L and O show thrice magnified sections of the regions enclosed in white boxes. The area enclosed within the white dotted rectangles drawn over the dorso-lateral epidermis adjacent to the PAS-DME interface in panels K and N shows the region of measurement of Rho1 intensity which further plotted in the graph Q. measurements were made in 25 embryos of each genotype an a P value≤0.05 from paired T test was considered to be significant and represented by *, whereas a P value ≤ 0.01 was considered to be highly significant shown by **. White arrows in panel N shows regions of elevated Rho-1 expression in conditional *Rab11* mutants.

The ANCs along with Cadherin-catenin complexes are major hubs of Rho signaling (Priya et al, 2013), required for the maintenance of the integrity of apical domain of epithelial cells. Thus, the cytoplasmic accumulation actin complexes in *Rab11* LOF mutants could trigger a cytoplasmic Rho signaling, as Rho signaling is intimately associated with cytoskeletal and cell-cell adhesion dynamics (Ohashi et al, 2017) and to confirm this, stage 14 *Rab11* mutant embryos with DC defects were immunostained for Rho1 (Fig 6 J-O). It was observed that Rho1 expression levels were indeed elevated in the DLEs of *Rab11* mutants as compared to the controls (insets in Fig 7 L and O), supporting our hypothesis.

**Fig. 7.**
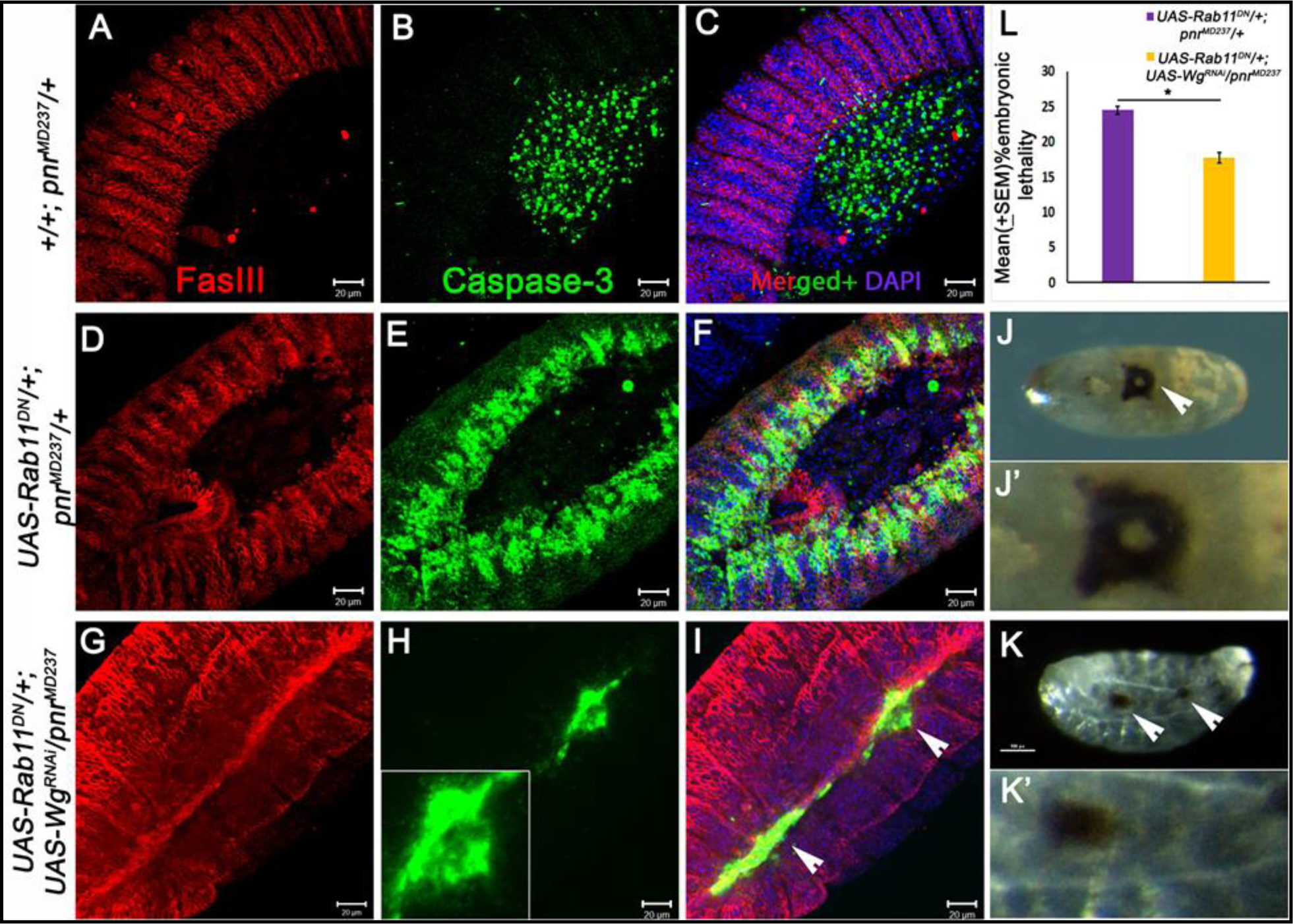
Functional Loss of Rab11 induces Caspase-3 expression in the DLEs which is rescued by a parallel Wingless knockdown. (A-I) Confocal projection images of stage 13 -14 developed embryos immunostained for FasIII (Red) and Cleaved Caspase 3(Green) and counterstained with DAPI (Blue) where (A-C) represents the undriven embryo, D-F represent projection images of *pnr-Gal4* driven *UAS-Rab11^N124I^*embryo where Caspase -3 shows a strong expression in the DLEs as compared to the undriven controls and panels G-I represent the *pnr- Gal4* driven *UAS-Rab11^N124I^;UAS-Wg^RNAi^*embryos which show a significant lowering of Caspase expression shown in white arrows in panel I. Elevated Caspase expression is observed only in regions which fail to zipper after the completion of dorsal closure process shown in the inset of panel H. *pnr-Gal4* driven *UAS-Rab11^N124I^*embryos not only shows a dorsal opening but also a significant amount of necrosis around the dorsal hole (J-J’), which is significantly rescued in the *pnr-Gal4* driven *UAS-Rab11^N124I^; UAS-Wg^RNAi^* embryos as they hatch out as first instar larvae although the dorsal lesions persist(shown in white arrows). This rescue is also confirmed through the embryonic lethality data represented by the graph L where a P value ≤ 0.05% was considered to be significant.

### *Rab11* shows a genetic interaction with *wingless* while regulating Caspase activation in the DLEs of embryos undergoing dorsal closure

Armadillo, in the absence of functional Rab11, accumulates within the cytoplasm of the DLEs and also elevates Rho1 expression in the DLEs of the *Rab11* mutant embryos which could be a result of the cytoplasmic accumulation of ANCs which remain stabilized at the Adherens junctions. Here we propose that an elevated expression of Armadillo and Rho1 in *Rab11* LOF mutants could mimic an elevated Wingless signaling condition if not up-regulate actual Wingless expression. It was observed that the mean % lethality of 24.48 ± 0.54% obtained in the embryos wherein *Rab11* was conditionally knocked out, was reduced to 17.71 ± 0.75% in embryos wherein Wingless was simultaneously knocked down (Table T4, Fig 7 L). It was also observed that approximately all 98.53±0.87% (data not shown) Rab11 knock out embryos showed an elevated expression of Caspase-3 in their DLEs (Fig 7 E and F) during DC, whereas only 67.87+0.76% (data not shown) of *pnr-Gal4* driven *UAS-Rab11^N124I^/+; UAS-Wg^RNAi^* embryos showed this expression, however, 26.98 +0.56% (data not shown) of *pnr-Gal4* driven *UAS- Rab11^N124I^/+; UAS-Wg^RNAi^* embryos showed a significant decline of Caspase expression in the DLEs with a partial accomplishment of DC. A partial Caspase expression was evident along the dorsal midline of the embryos (inset in Fig 7H) along with partial lesions (Fig. 7 K-K’). The first instars do not molt into the second instar stage suggesting a partial rescue of the *Rab11^DN^* phenotype as seen in Fig 7 J-J’. In order to confirm the interaction of *wingless* with *Rab11* in DC, *UAS-Rab11^RNAi^; UAS-Wg^RNAi^* and *UAS-Rab11^RNAi^* transgenic flies were driven with *pnr-Gal4* and a similar rescue in embryonic lethality was observed (Supplementary Fig. S3). This suggests that *Rab11* could potentially down-regulate Wingless signaling and consequently could prevent a probable JNK mediated apoptosis in the DLEs of fly embryos undergoing DC.

### *Rab11* maintains a proper JNK-Dpp signaling in the DLEs of the developing embryos through its genetic interaction with *wingless*

The JNK–Dpp signaling during dorsal closure regulates properties of DLEs which enable them to elongate and extend over the dorsal opening (Noselli and Agnes, 1999, Sluss and Davis, 1997). Reports of Thomas et al, 2009 and Tiwari and Roy, 2009 suggest the interaction of Rab11 and Rab30 with the JNK pathway during morphogenesis. This intrigued us to look into the effects of *Rab11* mutations in the JNK-Dpp expression pattern in the DLEs of stage 13-14 embryos, as, Rab11 also seems to regulate the expression of cadherin-catenin complex, which regulate JNK expression through the Egfr signaling. In order to monitor the Dpp expression pattern, *Rab11* mutant embryos with a Dpp-Lac-Z reporter were immunostained for β- Galactosidase where Dpp appeared to be significantly overexpressed (Fig 8C-C’). Similarly, to monitor the effects of *Rab11* LOF mutation on the JNK signaling pathway which plays an essential role during DC (Goberdhan and Wilson, 1998), a transgenic JNK biosensor stock, *TRE-dsRED/ TRE-JNK* (Chatterjee and Bohmann, 2012), duly introgressed with *pnr-Gal4* was used to drive, *UAS-Rab11^DN^* mutation where an elevated expression of RFP in the DLEs was fairly evident (Fig. 8 E-G). To further corroborate this hypothesis, stage 13-14 *pnr-Gal4* driven *UAS- Rab11^RNAi^* and *EP3017* homozygous embryos were also observed for JNK activity, where, it was found that *pnr-Gal4* driven *UAS-Rab11^RNAi^* showed a predominantly JNK down-regulated phenotype in the DLEs in 93.45±0.72% (data not shown) of the driven embryos (also reported by Nandy and Roy, 2020), however, a few embryos ∼5.35 +0.57% showed mild elevated expression towards the anterior end (shown with white arrows in Supplementary Figure S4-B’). 8.75 ±0.54 % (Data not shown) *EP3017* homozygous embryos on the other hand showed a fairly ubiquitous up-regulation of JNK signaling with more elevated expression of RFP near the dorsal regions (Supplementary Fig. S4-C’).

**Fig. 8.**
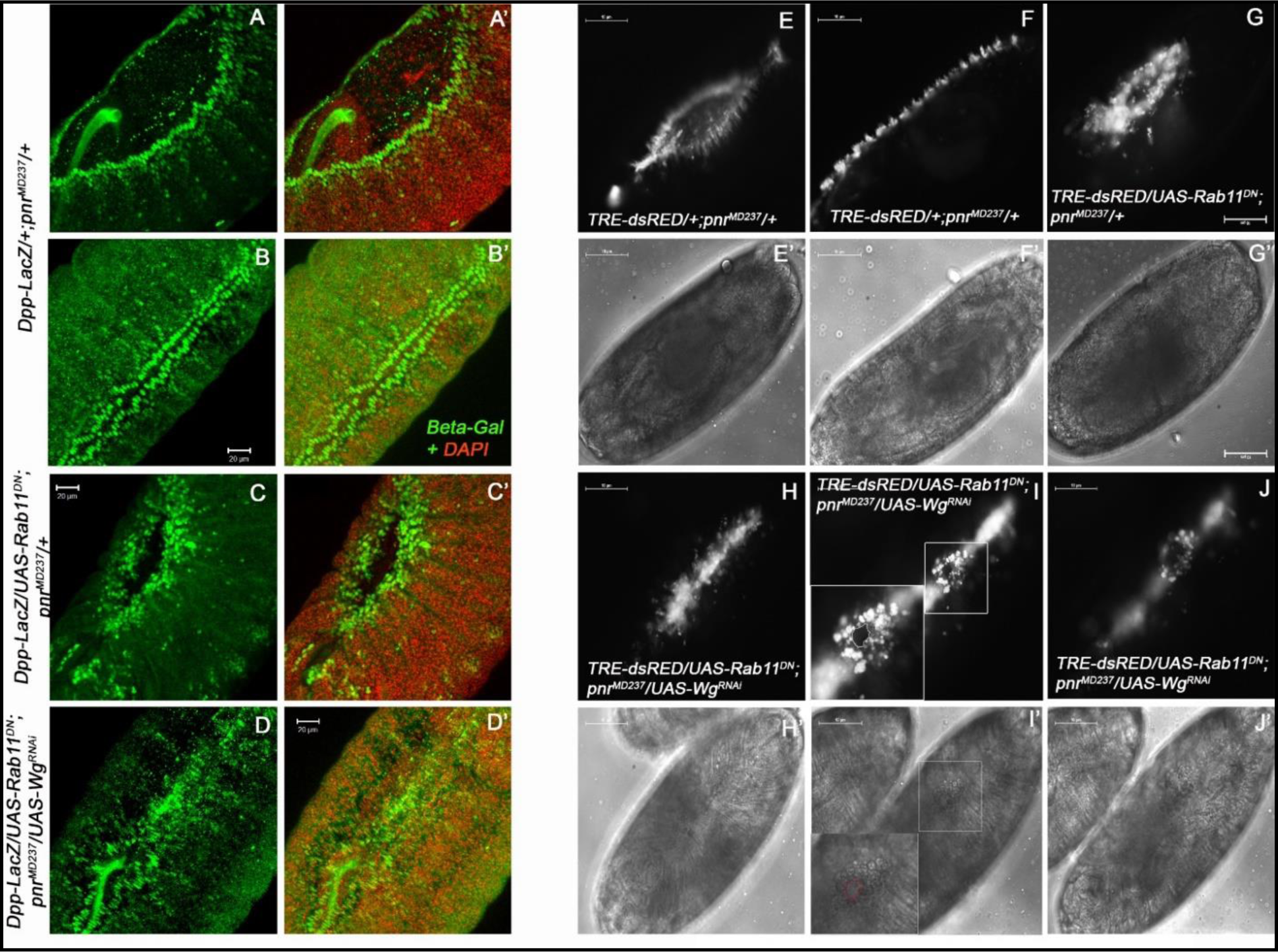
*Rab11* genetically interacts with *wingless* in the regulation of The JNK-Dpp pathway during dorsal closure. (A-B’) Confocal projection images of early stage 14 (A-A’) and stage 15 (B-B’) undriven embryo, immunostained for Beta-Galactosidase (Green) and DAPI (Red). (C-C’) represents a late stage 14, a *pnr-Gal4* driven *Dpp-LacZ/UAS-Rab11^N124I^*embryo where Dpp is elevated in the DLEs around the dorsal opening. D-D’ represents a stage 15 *pnr- Gal4* driven *dpp-LacZ/UAS-Rab11^N124I^; UAS-Wg^RNAi^*, with a partial restoration of the Dpp expression as compared to B-B’. In the embryo shown in D-D’, the regions along the dorsal axis where incomplete DC is clearly visible. (E-E’) and (F-F’) panels showing fluorescence and corresponding phase contrast images of stage 14 and stage 15 *TRE-dsRed; pnr-Gal4* embryos showing a robust expression of JNK in the DMEs. (G-G’) shows the fluorescence and the corresponding phase contrast image of *TRE-dsRED; pnr-Gal4* driven *UAS-Rab11^N124I^*embryo. Note the elevated expression of RFP around the dorsal hole of the embryo which suggests an increased JNK signaling.. H-H’, I-I’ and J-J’ show Fluorescence and Phase contrast images of *pnr-Gal4* driven *TRE-dsRED/UAS-Rab11^N124I^; UAS-Wg^RNAi^/+* rescued embryos, where in H-H’ there is a partial rescue in *pnr-Gal4* driven *UAS-Rab11^DN^* embryo. I-I’ and J-J’ are the images of the same embryo s where inset in I shows a partial dorsal opening which increases with time as shown in J which could result into a dorsal lesion as seen in the rescued embryos.

As reported by Kolahgar et al, 2011, the integrity of apical domain is essential for the prevention of JNK mediated apoptosis in the embryonic epithelial cells which is sincerely maintained by *Rab11* functions, hence, a loss of *Rab11* function in the dorsal cells results in the elevation of JNK expression which could ultimately elevate the expression of its downstream targets such as MMP1, Dpp and Caspase-3, as observed in our experiments. In this context, we were also intrigued to look into the interaction of *Rab11* with *wingless* in the regulation of JNK signaling. In agreement to our expectation, a rescue in the elevated JNK expression pattern (observed in *pnr-Gal4* driven *UAS-Rab11^DN^* embryos) was observed in 25.67±0.76% of *pnr-Gal4* driven *UAS-Rab11^N124I^; UAS-Wg^RNAi^* embryos, much like the undriven controls (Fig 8 H-J). Similarly, 23.34±0.45% of *pnr-Gal4* driven *UAS-Rab11^N124I^; UAS-wg^RNAi^* embryos showed a rescue of the Dpp phenotype, observed in the conditional *Rab11*-LOF mutants (Fig.8 D-D’).

## Discussion

Coordinated cell shape changes in the fly embryonic epithelium during morphogenesis and dorsal closure requires the strict retention of epithelial properties of cells, such as cell polarity, cell-cell/matrix adhesions, and cytoskeletal organization, contrary to the single cell shape changes (Plastushenko and Blanpain, 2019). Thus maintenance of epithelial integrity (in terms of cell polarity and cell-cell adhesions) at the time of shape modification is a must for the DLEs during dorsal closure as they elongate in a coordinated manner over the dorsal opening. Recycling Endosome being a significant contributor of the membrane and cytoskeleton organization of epithelial cells (Welz et al, 2014), therefore, seems to be showing promising roles in DC, especially when coordinated cell shape changes of the DLEs are concerned. Earlier experiments have revealed that a constant Rab11 expression in the fly embryonic epithelium is a must for the maintenance of cell-cell/matrix adhesions such as E-Cadherin (Choubey et al, 2020), Integrin (Bhuin and Roy, 2012) and septate junction organization, along with cortical actin organization and cell polarity dynamics (Nandy and Roy, 2020). Experiments of Harden et al, 1999 have demonstrated the involvement of small Ras like GTPAses such as Rho1, dRac and cdc-42 in the regulation of the core JNK-Dpp signaling in the DLEs which is essential for dorsal closure (Noselli and Agnes, 1999 and Stronach and Perrimon, 2002). Therefore, it becomes important to address the DLE specific functions of Rab11 during dorsal closure.

In order to observe the effects of a complete knock-out of *Rab11* from the DLEs, The *Rab11^N124I^* allele was considered ideal, as it specifically targets and inhibits Rab11 function by binding to the GTPase domain of active Rab11 (Duman et al, 1999 and Satoh et al, 2005). Surprisingly, not only DC defects were evident in all the *pnr-Gal4* driven *UAS-Rab11^N124I^* embryos, the cells of the DMEs underwent apoptosis (Fig 3 and Supplementary Fig S2) suggesting a probable role of *Rab11* in the negative regulation of Caspases in the DLEs during DC. Several reports suggest that the activation and expression of Caspases in the amnioserosa cells which help in the generation of forces, required to contra-laterally drag the lateral epithelia over the dorsal opening (Teng et al, 2011; Sokolow et al, 2012; Muliyil et al, 2011; Toyama et al, 2008). Our experiments suggest a parallel necessity of Caspase downregulation in the DMEs at the same time which is perhaps related to Rab11 functions.

Apoptosis in an epithelium may be triggered by the loss of a pre-defined, differentiated state of the constituent epithelial cells which includes parameters like epithelial cell membrane organization, cytoskeletal organization and cell polarity (Jesowzka et al, 2011; Kolahgar et al, 2011). The cortical actin and Myosin V organization suffers a substantial loss in *Rab11* LOF mutants (Fig. 4) suggesting cytoskeletal disruption which could be a prime cause of the inability of the respective cells to undergo extensions in order successfully complete the dorsal closure process.

The cortical actin filaments remain anchored to the apically located adherens junctions of epithelial cells as reported by Tsukita et al, 1999; Sawyer et al, 2009; Kametani and Takeichi, 2007, Borghi et al, 2012, and its role in epithelial morphogenesis and tissue remodeling is an indispensable one (Niessen et al, 2011 and Tepass, 1999). The fact that Cadherins and cortical actin filaments get disrupted in a *Rab11* mutant background (Desclozeaux et al, 2008), could result in an ectopic and untimely manifestation of EMT which was confirmed from the elevated expression of MMP1 in *Rab11* mutant embryos. Interestingly, Cadherins communicate to the cortical cytoskeleton through anchorage molecules like the catenins.

The α and β Catenins are responsible for the intracellular transduction of biophysical signals from membrane associated Cadherins (Gumbiner and McCrea, 1993). Here Armadillo/ β-catenin seems to play an instrumental role as it binds to the actin associated α-Catenin to the membrane associated Cadherin (Pai et al, 1996). The increase of intracellular Armadillo in *Rab11* mutants could be a direct consequence of its dissociation from the Adherens junction complex which could be due to the proteolytic cleavage of Cadherin associated Armadillo by Caspase-3 as suggested from their co-localization. The results also indicate that the organization of Armadillo at the Cadherin-catenin complex may not be only regulated by Rab11 and Armadillo accumulation in the DLEs could be a Caspase mediated effect. Interestingly, the cytoplasmic accumulation of Armadillo in Rab11 mutant DLEs is unique as perhaps it replicates an active Wingless signaling condition.

The Wingless pathway plays an instrumental role in the process of dorsal closure (Morel and Arias, 2004 and McEwen et al, 2000), where the McEwen group elucidates the genetic interaction of the Wingless pathway with the core JNK-Dpp pathway in the process of dorsal closure. This suggests that a Wingless elevation could probably result in an elevated JNK signaling as observed in *Rab11* mutants (Fig. 5). Again the elevated expression of Rho1 observed in the DLEs of *Rab11* mutant embryos could retard the differentiation process of the DMEs as they elongate and extend during DC, where reports of Hariharan et al, 1995, show that Rho1 overexpression exerted deleterious effects in developing *Drosophila* eyes. Also, the possibility of Rho1 expression in the DLEs, through the non-canonical Wingless signaling pathway cannot be ignored (Winter et al, 2001). We observed that a simultaneous knockdown of Wingless in *Rab11* LOF mutants i.e. in *pnr-Gal4* driven *UAS-Rab11^N124I^; UAS-Wg^RNAi^* embryos, not only a resulted in a decrease of embryonic lethality, but also in the decrease of the severity of the necrotic patch observed in *pnr-Gal4* driven *UAS-Rab11^N124I^* embryos, enabling them to reach up to the 1^st^ instar stage of larval development (Fig. 7) establishing the genetic interaction of *wingless* with *Rab11* in the regulation of Caspase expression in the DMEs during DC.

Several reports suggest that the JNK pathway is critically important for cellular motility and epithelial morphogenesis (Xia and Karin, 2004), and leads to the downstream expression of genes such as *dpp* (Sluss and Davis, 1997; Riesgo-Escovar and Hafen, 1997), *puckered* (Martin Blanco et al, 2000) *mmp1* (Jin et al, 2011) *integrins* (Homsy et al, 2006) and *Caspases* (Kolahgar et al, 2011). The up-regulated expression of MMP1 along with Caspase-3 in *Rab11* mutants could be a consequence of an elevated JNK expression as seen in Fig. 8 G. Interestingly, Dpp expression in DLEs of *Rab11* mutants was elevated too (Fig 8 A-C’) which could be a consequence of elevated JNK levels. Dpp, although elevated in *pnr-Gal4* driven *UAS-Rab11^DN^* mutants could not perhaps perform its conventional function of cell transformation or cell survival, probably, due to the inability of Rab11 to recycle membrane associated Tkv (Deshpande et al, 2016). Also, in a *Rab11* LOF condition, Dpp may fail to be exocytosed (Entchev and Gonzalez Gaitan, 2002). Considering this explanation we further believe here that Rho1 overexpression may not be Dpp induced, instead, could be either a consequence of a disrupted PCP program or may be directly induced by a *Rab11* LOF condition which could prevent ERM (Ezrin, Radixin and Moesin) adhesion assembly with a consequent expression of cytoplasmic Rho1 followed by a Rho1 induced JNK mediated apoptosis (Neisch et al, 2010) in the DLEs. However, this hypothesis requires further experiments and validation which is beyond the scope of this report.

Our experiments demonstrate a functional relevance of Rab11 in the physiology of DLEs at the time of *Drosophila* embryonic dorsal closure, an ideal model to study the cellular processes involved in dorsal closure. We show here that the precisely controlled expression of JNK in the DLEs of fly embryos is Rab11 dependent, which if lost may result in the elicitation of an apoptotic program. Here, apart from Rab11’s conventionally reported roles of apical recycling and maintenance of JNK signaling, we also show that it may regulate the intracellular pools of membrane associated molecules, a perturbation of which may mimic an overexpression condition of a critically important signaling pathway such as Wingless. Thus we suggest it would be extremely interesting to deduce and confirm the afore-mentioned mechanism involving Rab11 functions in higher systems such as neural tube closure (Ossipova et al, 2014), vertebrate gastrulation (Ossipova et al, 2015) and cancer metastasis (Xu et al, 2016).

## Materials and Methods

### Fly stocks and rearing conditions

Fly stocks have been reared on standard food preparation containing maize powder, agar, yeast and sugar and methyl-p-hydroxy benzoate was added as anti-fungal along with propionic acid as anti-bacterial agents at a temperature of 23±1°C. The stock *pnr^MD237^/TM3, Ser* was obtained from Bloomington *Drosophila* Stock Center (BDSC 3039, Thomas et al, 2009) and expresses *Gal4* as reported by Calleja et al, 1996. This stock was further introgressed with *TM3, Act-GFP, Ser ^1^/TM6B* in order to generate *pnr^MD237^/TM3, Act-GFP, Ser^1^*stock. *TRE-JNK* (Chatterjee and Bohmann, 2012) was introgressed with *Sp/CyO; pnr^MD237^/ TM3, Act-GFP, Ser^1^* and *Sp/CyO; pnr^MD237^/TM6B* to obtain *TRE-JNK/CyO*; *pnr^MD237^/TM3, Act-GFP, Ser^1^* stock, respectively.*UAS- Rab11^N124I^ (UAS-Rab11^DN^)*/*CyO* stock (Satoh et al, 2005 a kind gift from Dr. Ready) was introgressed with *Tft/CyO-Act-GFP* stock in order to generate *UAS-Rab11^N124I^/CyO-Act-GFP* stock*, EP3017/TM6B* () was introgressed with *TM3, Act-GFP, Ser^1^/TM6B* to generate *EP3017/TM3-Act-GFP,Ser1* stock which was further introgressed with *TRE-JNK/CyO-Act-GFP; Dr/TM3-Act-GFP* stock to generate *TRE-JNK/CyO-Act-GFP; EP3017/TM3-Act-GFP*, *Ser^1^* stock. *UAS-Wg^RNAi^* (BDSC 32994) was double balanced to generate *Sp/CyO-ActGFP; UAS- Wg^RNAi^* stock which was further used to generate *UAS-Rab11^N124I^/CyO-Act-GFP; UAS-Wg^RNAi^* stock*. UAS-p35* (BDSC 5073) stock was double balanced to generate *Sp/CyO-Act-GFP; UAS- p35* stock which was further introgressed with *UAS-Rab11^N124I^/CyO* stock to generate *UAS-*

*Rab11^N124I^/CyO-Act-GFP; UAS-p35* stock. *Dpp-lacZ/CyO* (BDSC 68153) flies were double balanced to obtain *Dpp-LacZ/CyO-Act-GFP; dco^2^/TM3, Act-GFP, Ser^1^* stock which was further introgressed with *Sp/CyO-Act-GFP; pnr^MD237^/TM3, Act-GFP, Ser^1^* to obtain *Dpp-LacZ/CyO, ActGFP; pnr^MD237^/ TM3, Act-GFP, Ser^1^* stock. This stock was used as a driver in our experiments where only non-GFP embryos obtained from the crosses set with the flies containing *UAS* transgenes were selected and then proceeded for β-Galactosidase staining.

With the purpose to observe the tissue specific effects of a targeted expression *Rab11* LOF mutation in the dorso-lateral epithelial cells of DC undergoing embryos, the *Gal4-UAS* system of targeted or conditional gene expression as described by Brand and Perrimon, 1993, was used to drive its dominant negative allele in stage13 and 14 fly embryos. *pnr^MD237^/ TM3, Act-GFP, Ser^1^* stock was used as the *Gal4* to drive different mutants under the *UAS* promoter in order to observe DLE specific effects. Males of *Gal4* stock and Virgin Females of *UAS* transgene containing lines were used to set up crosses in order to obtain embryos of the desired genotype. In order to ensure the proper segregation of the transgenic *Gal4* and *UAS* constructs in an individual, the transgene bearing parental stocks were balanced with green balancers and the progeny obtained from their introgression were screened for the absence of the GFP containing balancer chromosomes. The balancers containing Act-GFP construct showed the presence of the actin rich haemocytes under the fluorescence microscope which were completely absent in the embryos devoid of the balancers (Fig. 1 C-E’). These embryos were further staged as suggested by Green et al, 1993. The embryos of identical stages were further used for analysis of cell-biological and molecular parameters.

### Embryo collection and fixation

Flies were made to lay eggs on standard agar plates supplemented with sugar and propanoic acid and the egg collection was done as per the protocol suggested by Narasimha and Brown, 2006, with slight modifications. For making whole mount preparations and immunostaining, different alleles and transgenes were balanced with GFP tagged balancers and only non GFP or driven embryos were selected for experimental purpose. Eversed clypaeolabrum was treated as a marker of stage 13 and retracted clypaeolabrum was treated as marker of stage 14 (Sasikumar and Roy, 2009). Also, 1 h synchronized embryos were chosen for immunostaining experiments. Embryo staging was done according to Hartenstein’s Atlas of *Drosophila* Development, 1993.

### Genetics

#### Embryonic cuticle preparations

Embryonic cuticles were prepared as described by Anderson, 1985; Ostrowski et al, 2002 with slight modifications. (Nandy and Roy, 2020).

#### Immunostaining, imaging and confocal microscopy

Fixation and imaging of *Drosophila* embryos was done as described by Narasimha and Brown, 2006. Embryos once dechorionated and devitellized were fixed in 4% para-formaldehyde solution and stored in absolute methanol. For immunostaining, these embryos were rehydrated using descending methanol gradients of 70%, 50%, 30% and 10% in 0.1% PBT solution. The embryos were blocked for 2h at RT in blocking solution as described by Banerjee and Roy, 2017. Rabbit anti-sera against *Drosophila* Rab11 (Alone et al, 2005) was used at a dilution of 1:100 for immunostaining and 1:1000 for western, anti-FasIII (7G10, DSHB), FITC conjugated mouse anti-p-Tyr antibody (Sigma-USA) used at a dilution of 1:100, anti-Caspase-3 antibody (Sigma USA, Dubey and Tapadia, 2017) was used at a dilution of 1:100, anti-Hid antibody (Santa Cruz) used at a dilution of 1:20, anti MyoV (A kind gift from Dr. A. Satoh of Hiroshima Unversity (Japan), Satoh et al, 2005) used at a dilution of 1:200, anti- Armadillo (N27A1, DSHB) used at a dilution of 1:100, anti Rho-1(p1D9, DSHB) used at a dilution of 1:50, anti MMP1(5H7B11, DSHB) used at a dilution of 1:100 and anti-β-Galactosidase antibody (Invitrogen) raised in chicken was used at a dilution of 1:500. Secondary antibodies were used as described by Sasikumar and Roy, 2009; Bhuin and Roy 2012; Ray and Lakhotia, 2018, and were imaged using single photon confocal microscope using Zen software, 2012. The images obtained were analyzed using Zeiss LSM-510 Meta-software (Including aspect ratio measurement Fig.2). Quantification of fluorescence intensities was done by the Adobe Photoshop CS6 software where regions of intensity measurement were selected under the “lasso” or “rectangular marquee” tool and values were obtained from the “Histo” tool. Dark field, fluorescence and phase contrast images of the embryos were taken under the Nikon eclipse E800 microscope under the same gain and exposure values. 9-11h developed embryos were de-chorionized and mounted in halocarbon oil in order to image them live in phase contrast as well as fluorescence microscope.

#### Embryonic, pupal and larval lethality assays

The *Gal4-UAS* system of targeted gene expression was used in order to see the effects of alleles of *Rab11* on the embryonic lethality, wherein every experiment males of the genotype *pnr- Gal4/+* and virgin females with transgenic *UAS* constructs, i.e., *UAS-Rab11^N124I^*, *UAS- Rab11^N124I^; UAS-wg^RNAi^* etc. were introgressed and embryos were collected from the F1 generation (balancers were removed from the parental stocks in order to nullify their effects). These embryos were incubated for 24 to 48 h at 23°C on standard agar plates and the total number of dead embryos (detected by yellowing of the eggs) was counted against total number of fertilized eggs. These fertilized eggs include the dead as well as the hatched embryos, and percentage death was calculated as:

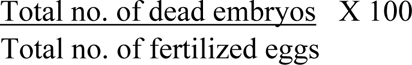

The % lethality for each cross was calculated in triplicates with a total number of 200 embryos in each set of observation, thus a gross total of 600 embryos were observed. The mean lethality observed in each case was tabulated and graphically represented along with standard deviation and standard error values. Paired T-test was done to determine the significance between the lethality values of two comparable genotypes where a p value ≤ 0.05 was considered as significant.

#### Calculation of mean aspect ratios

In order to assess the extent of shape changes induced by the *Rab11^DN^* mutation we calculated the mean aspect ratio of the DMEs and LEs in the *Rab11* LOF condition as well as in the undriven controls where maximum length and maximum breadth of 50 individual cells (from separate embryos) of DMEs and LEs were measured using the LSM-510 Meta software and then maximum length: maximum breadth ratio was calculated for each cell. These values were then graphically plotted using the MS-Excel-Worksheet software.

## Acknowledgements

We thank the fly community for generously providing fly stocks. Special acknowledgements to Professor B.J. Rao for providing the TRE-JNK/CyO fly stock and Dr. A.K. Satoh for providing the anti-Myo V antibody. We thank Dr. Satish Sasikumar and Dr. Tanmay Bhuin for their pioneering studies on the functions of Rab11 on *Drosophila* embryonic epithelial morphogenesis. Special thanks to Mr. Rohit Kunar for his critical suggestions during the preparation of the manuscript. We duly acknowledge the National facility for Laser Scanning Confocal Microscopy, Department of Zoology, Banaras Hindu University. The equipment support of DST-FIST and CAS Zoology are duly acknowledged.

## Author Contributions

Conceptualization: NN Methodology: NN Investigation: NN

Visualization: NN

Project administration: JKR Supervision: JKR

Writing – original draft: NN

Writing – review & editing: NN and JKR

## Data and materials availability

For obtaining any data from the manuscript, one may communicate with the corresponding author. The fly stocks and reagents used are available with the corresponding author as well as the resources which have been mentioned in the materials and methods section of the manuscript.

## Supplementary Data

**Supplementary Fig. S1.**
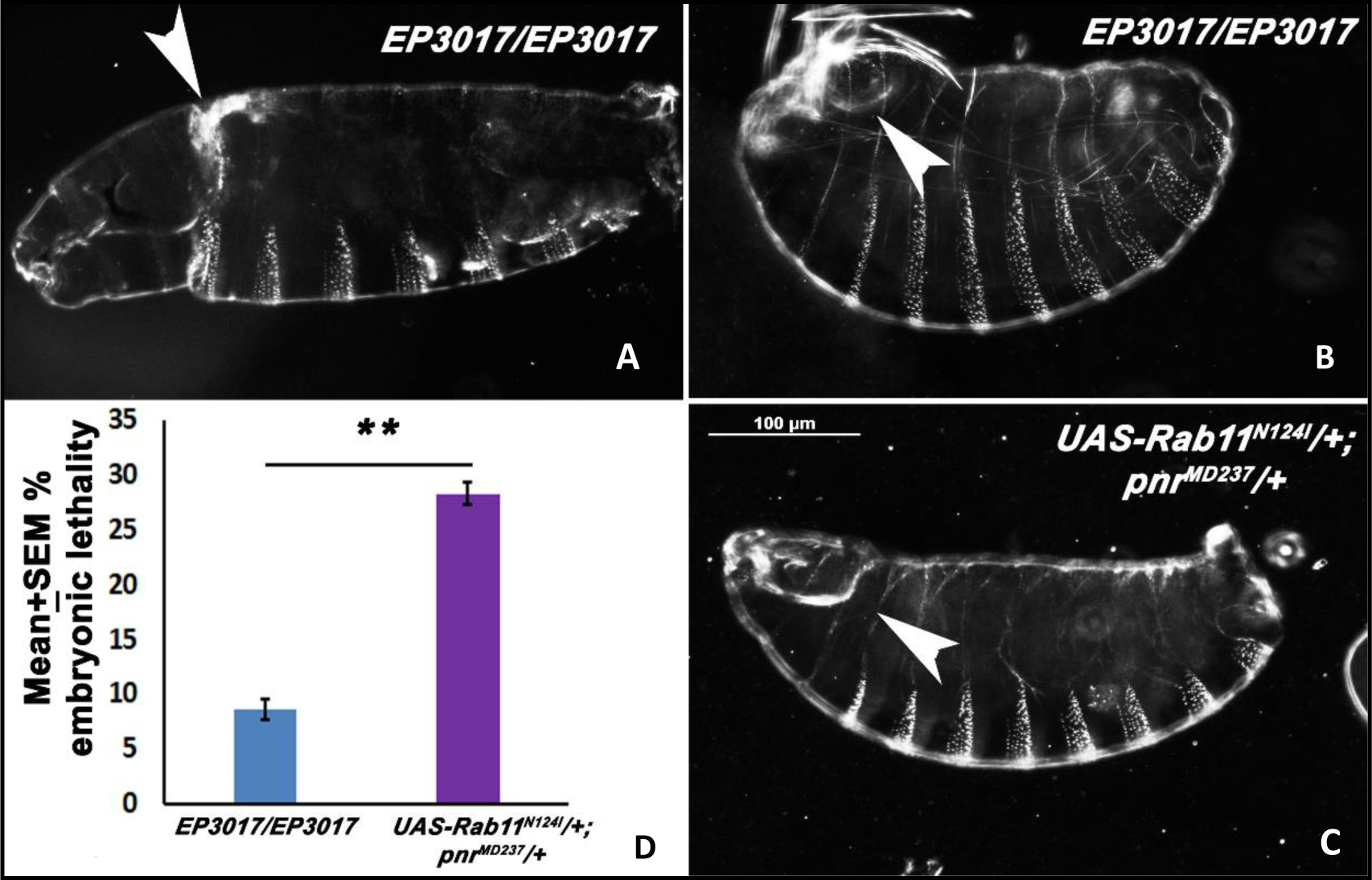
(in support of Fig 1): Dark field images of cuticles of *EP3017/EP3017* homozygous, first instar larva (A), 22-24 h developed embryo (B) and a *pnr-Gal4* driven *UAS- Rab11^N124I^* embryo (C). The anterior dorsal opening is quite conspicuous in the *EP3017* homozygotes which phenocopies the *pnr-Gal4* driven *UAS-Rab11^N124I^*embryos.At the same time there is a considerable shrinkage of the embryos along the anterior posterior axis, a characteristic ventral puckering and anterior dorsal opening much like the *EP3017* homozygous embryos. D- Graph representing the embryonic lethality% in *EP3017* homozygotes and *pnr-Gal4* driven *UAS-Rab11^N124I^*embryos. The lethality% observed in *pnr-Gal4* driven *UAS-Rab11^DN^* condition is significantly more as compared to *EP3017* mutants where 600 embryos of each genotype was observed and a P value ≤ 0.01 was considered as highly significant and represented by **.

**Supplementary Fig. S2.**
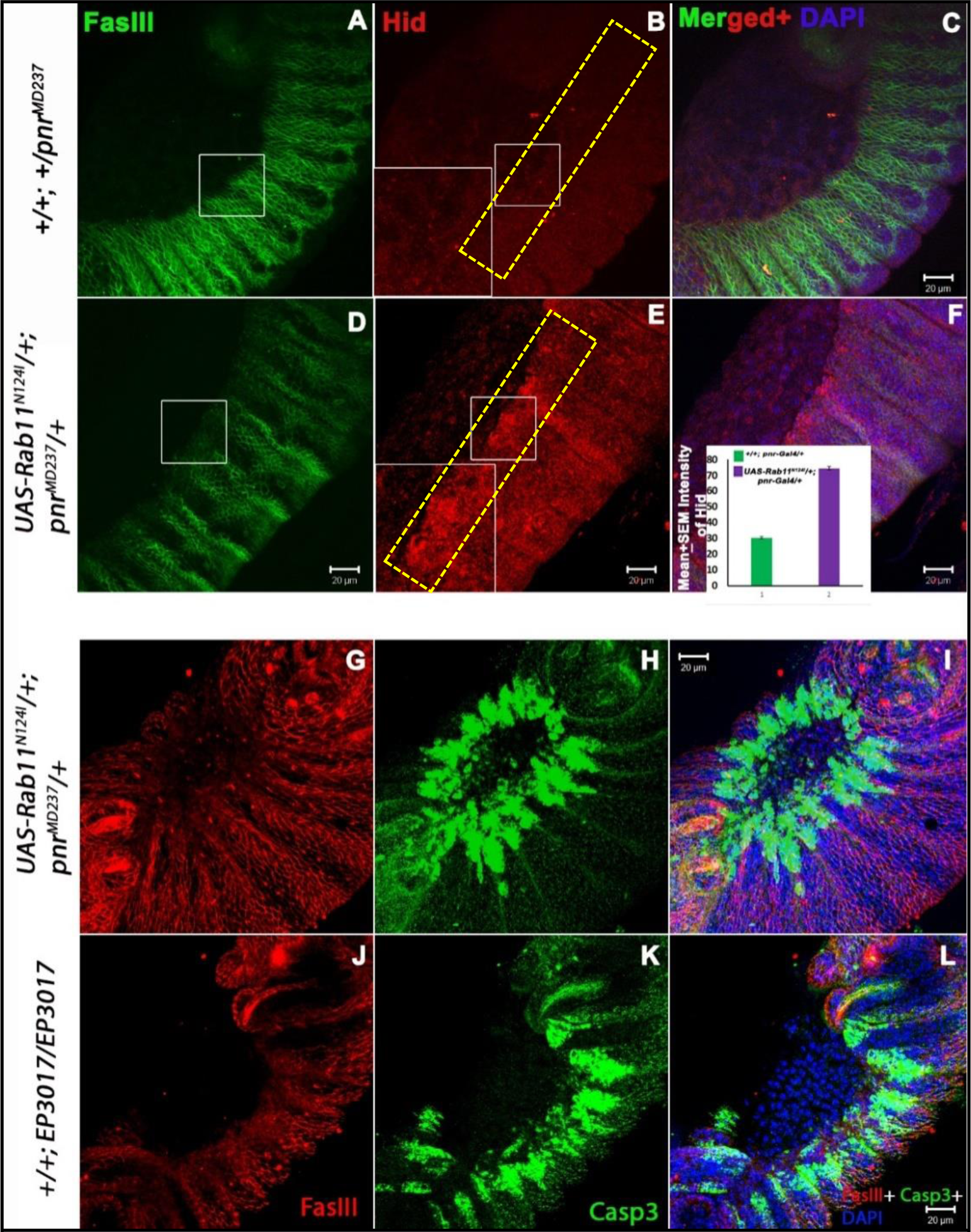
(in support of Fig 3): (A-F) Confocal projection images of stage 13 -14 embryos immunostained for FasIII (Green), Hid (Red) and counterstained for DAPI (Blue), where panels A-C represent the un-driven embryos and D-F represent the *pnr-Gal4* driven *UAS-Rab11^N124I^*embryos where improper cell morphologies are evident as shown in the white box in D. These cells show an elevated expression of Hid unlike the controls which has been measured in the area enclosed by the rectangles in B and E and graphically plotted in the panel F. The significant increase in the intensity Red fluorescence indicates an elevation of Hid expression in *pnr-Gal4* driven *UAS- Rab11^N124I^*mutants suggesting apoptosis in these cells.(G-L) represents the confocal projection images of embryos co-immunostained for FasIII (Red) and Caspase-3 (Green) and counterstained with DAPI (blue) where panels G-I represent *pnr-Gal4* driven *UAS-Rab11^N124I^* embryos where a surge of Caspase-3 expression is evident in the dorsolateral epidermis where there is a considerable loss of cell shape and cell membrane organization. (J-L) Panels representing *EP3017/EP3017* embryos which show considerable dorsal closure defects and also a surge in Caspase expression much like the *pnr-Gal4* driven *UAS-Rab11^N124I^* mutants (n=25 embryos in each case).

**Supplementary Fig. S3.**
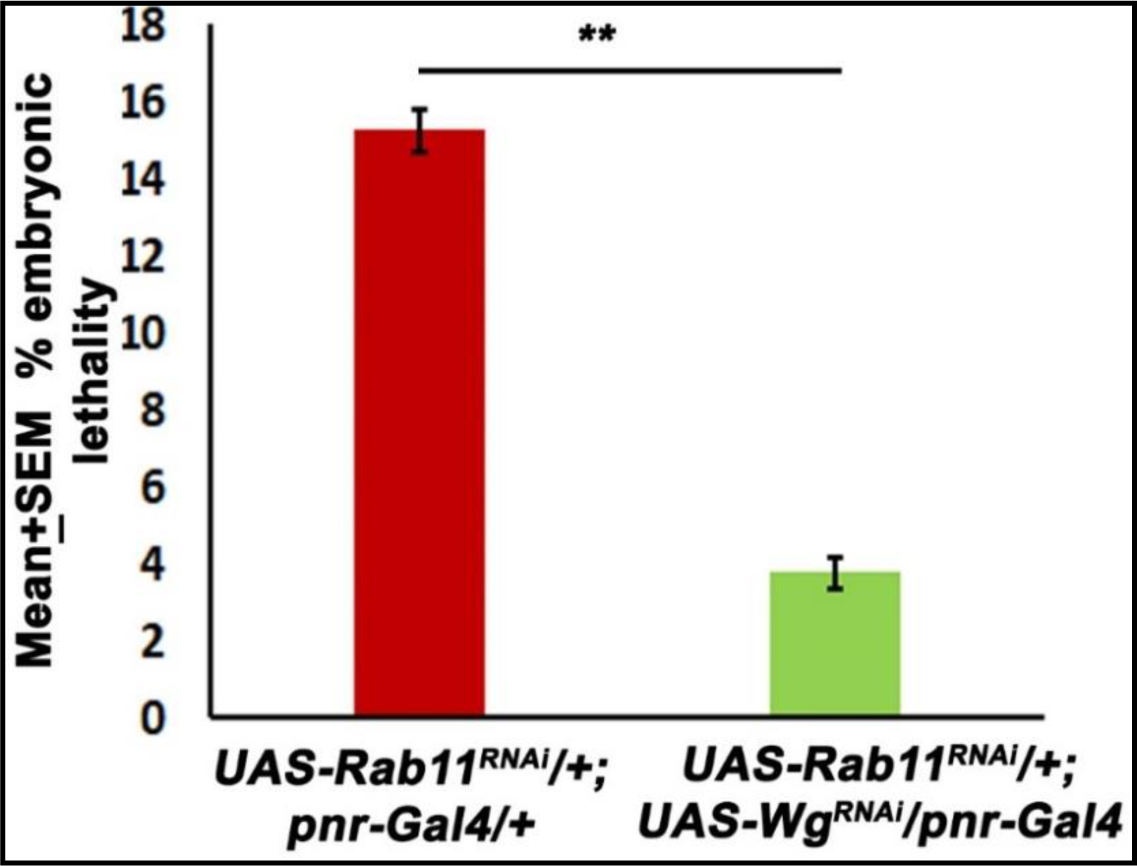
(In support of Fig. 7):Graphical representation of Mean ±% lethality of 22-24 h developed *Drosophila* embryos where *pnr-Gal4* driven *UAS-Rab11^RNAi^* embryos show an embryonic lethality of 15.28 ± 0.57% (Table T6), shown in the red bar whereas in *pnr-Gal4* driven *UAS-Rab11^RNAi^*; *UAS*-*wg^RNAi^* embryos a lethality of 3.75±0.39% was observed (shown in the green bar) which suggests a rescue of the dorsal closure defects caused due to a targeted knock-down of Rab11 in the dorsolateral epidermis with a simultaneous knock-down of *wingless*.

**Supplementary Fig.S4.**
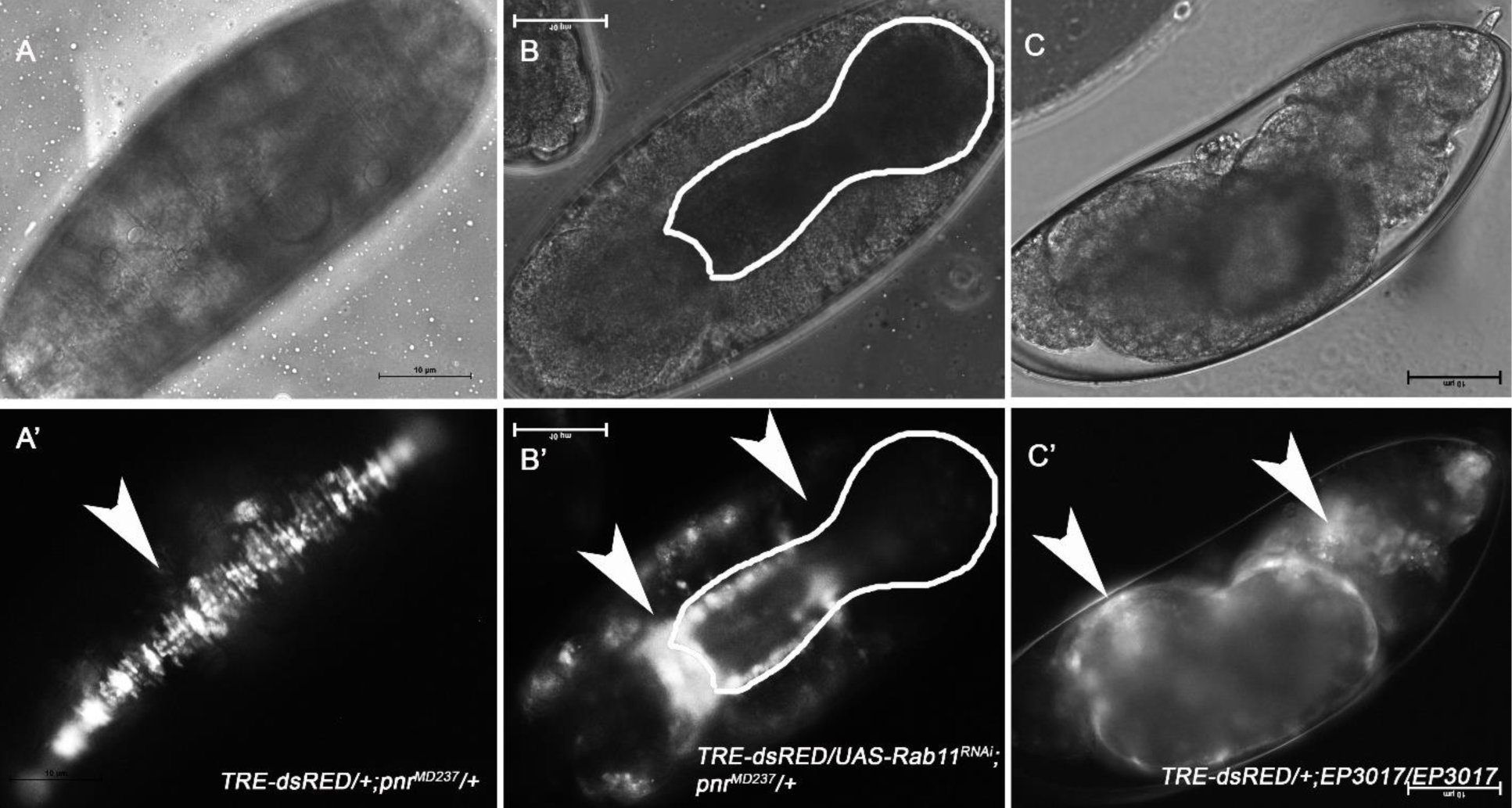
(in support of Fig.8): Phase contrast and Fluorescence images ofstage 13-14(A-A’) *TRE-dsRED/+; pnr^MD237^/+;* (B-B’) stage 13-14 *TRE-dsRED/UAS-Rab11^RNAi^; pnr^MD237^/+* and C-C’ *TRE-dsRED/+; EP3017/EP3017* embryos suggesting the expression pattern of JNK in the DLEs of the embryos which are undergoing dorsal closure. White arrow in A’ represents the robust expression of p-JNK in the DLEs of control embryos undergoing the dorsal closure process whereas on the other hand white arrows in panel B’ represents regions of elevated and downregulated JNK expression. White arrows in C’ represents regions of elevated JNK expression in the dorsal epidermis of the embryos with dorsal closure defective phenotypes. These evidences suggest a strong effect of *Rab11* LOF mutations on the JNK signaling pathway during dorsal closure.

**Supplementary Fig. S5.**
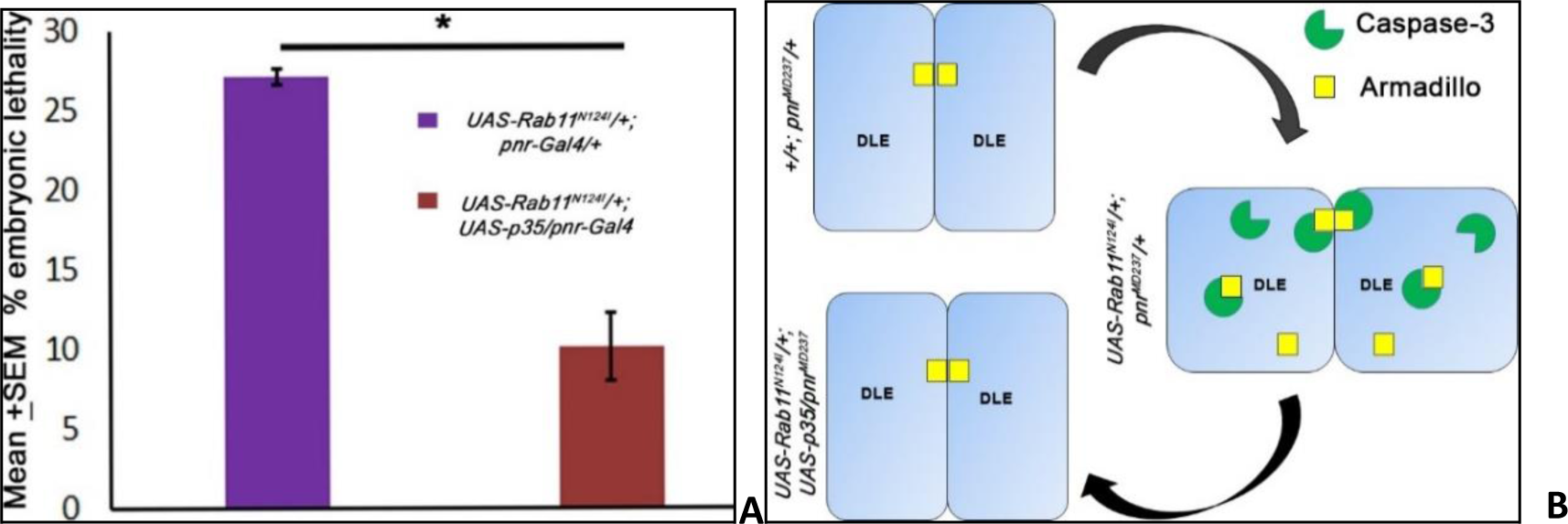
(In support of Fig. 5**): (A)** Graphical representation of Mean ±% lethality of 22-24 h developed *Drosophila* embryos where *pnr-Gal4* driven *UAS-Rab11^N124I^* embryos show an embryonic lethality of 27.1 ± 0.26% (Table T7), shown in the purple bar whereas in *pnr-Gal4* driven *UAS-Rab11^N124I^*; *UAS*-*p35* embryos a lethality of 10.08±1.21% was observed (shown in the crimson bar) which suggests a rescue of the dorsal closure defects caused due to a targeted knock-down of Rab11 in the dorsolateral epidermis with a simultaneous expression of the anti-apoptotic gene *p35*. **(B)** Schematic representation of spatial organization of Armadillo and Caspase-3 in the DLEs of fly embryos undergoing dorsal closure in undriven i.e. *pnr-Gal4/+, pnr-Gal4* driven *UAS-Rab11^N124I^* and *pnr-Gal4* driven *UAS-Rab11^N124I^; UAS-p35* genetic backgrounds drawn from the experimental evidences in Fig.5. The figure suggests a probable mechanism of intracellular Armadillo sequestration in conditional Rab11LOF mutants by a probable proteolytic action of Caspase 3 on the plasma membrane of the DLEs.

## Supplementary Tables

**Table T1.** (In support of Fig1):

|  | Total | unfertilized | fertilized | dead | Death% |
| --- | --- | --- | --- | --- | --- |
|  | 309 | 96 | 213 | 59 | 27.69953 |
| $UAS-Rab11^{N124I}/+; pnr-Gal4/+$ | 345 | 78 | 267 | 67 | 25.09363 |
|  | 323 | 87 | 236 | 52 | 22.0339 |
|  |  |  |  | Mean | 24.94235 |
|  |  |  |  | SD | 2.835844 |
|  |  |  |  | SEM | 1.637275 |
|  | 200 | 25 | 175 | 3 | 1.714286 |
| *+/+; pnr-Gal4/+* | 200 | 26 | 174 | 3 | 1.724138 |
|  | 200 | 25 | 175 | 2 | 1.142857 |
|  |  |  |  | **Mean** | 1.527094 |
|  |  |  |  | **SD** | 0.332795 |
|  | **P Value** | **0.003922091** |  | **SEM** | 0.192139 |

**Table T2.** (In support of Fig. 2):

|  |  | Mean AR | SD | SEM |  | Mean AR | SD | SEM |
| --- | --- | --- | --- | --- | --- | --- | --- | --- |
| *+/+; pnr-Gal4/+* | **DME** | 5.264189 | 1.712894 | 0.541665 | **LE** | 3.559061 | 0.693077 | 0.21917 |
| *UAS-Rab11<sup>N124I</sup>/+; pnr-Gal4/+* | **DME** | 2.07325 | 0.465438 | 0.147184 | **LE** | 1.602887 | 0.397438 | 0.125681 |

P Value for DME Aspect ratio: **6.63752E-05**
P Value for LE Aspect ratio: **5.18543E-05**

**Table T3.**
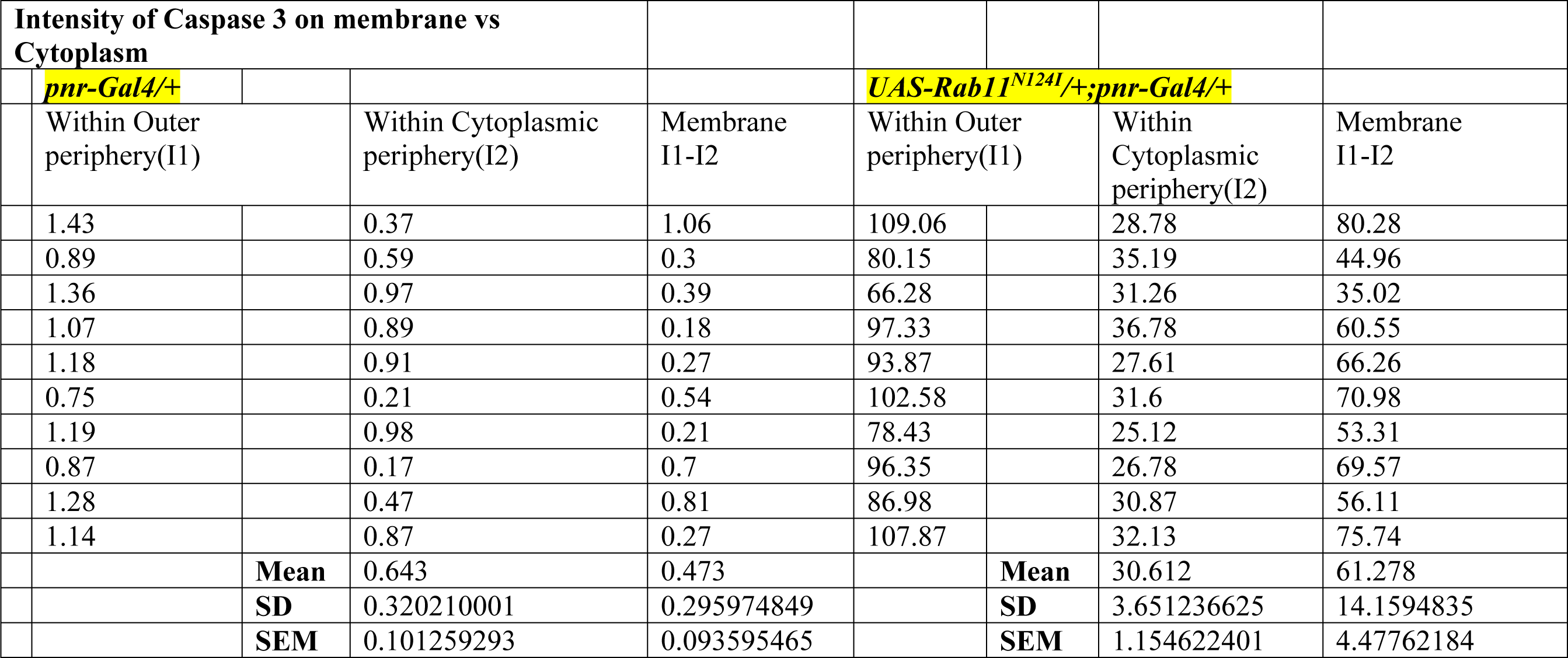
(In support of Fig.3)

**Table T4.** (In support of **Fig.8**):

| *UAS-Rab11<sup>N124I</sup>/+;*<br>*pnr-Gal4/+* |  |  |  |  | *UAS-Rab11<sup>N124I</sup>;*<br>*UAS-Wg<sup>RNAi</sup>/pnr-Gal4/+* |  |  |  |  |
| --- | --- | --- | --- | --- | --- | --- | --- | --- | --- |
| Total | Unfertilized | Fertilized | Dead | Death% | Total | Unfertilized | Fertilized | Dead | Death% |
| 300 | 9 | 291 | 74 | 25.42955 | 300 | 12 | 288 | 47 | 16.31944 |
| 300 | 11 | 289 | 68 | 23.52941 | 300 | 9 | 291 | 55 | 18.90034 |
| 300 | 10 | 290 | 71 | 24.48276 | 300 | 10 | 290 | 52 | 17.93103 |
|  |  |  | Mean | 24.48057 |  |  |  | Mean | 17.71694 |
|  |  |  | SD | 0.950073 |  |  |  | SD | 1.303701 |
|  |  |  | SEM | 0.548525 |  |  |  | SEM | 0.752692 |
|  |  |  |  |  | P<br>value | 0.017453236 |  |  |  |

**Table T5.** (In support of Supplementary Fig. S3):

|  |  |  |  |  |  |  |  |  |  |  |
| --- | --- | --- | --- | --- | --- | --- | --- | --- | --- | --- |
|  | *pnr-Gal4>>UAS-Rab11<sup>RNAi</sup>* |  |  |  |  | *pnr-Gal4>>UAS-Rab11<sup>RNAi</sup>;UAS-Wg<sup>RNAi</sup>* |  |  |  |  |
| Total | Unfertilized | Fertilized | Dead | Death% |  | Total | Unfertilized | Fertilized | Dead | Death% |
| 300 | 23 | 277 | 45 | 16.24549 |  | 300 | 39 | 261 | 8 | 3.065134 |
| 300 | 19 | 281 | 43 | 15.30249 |  | 300 | 52 | 248 | 11 | 4.435484 |
| 300 | 27 | 273 | 39 | 14.28571 |  | 300 | 33 | 267 | 10 | 3.745318 |
|  |  |  | Mean | 15.2779 |  |  |  |  |  | 3.748645 |
|  |  |  | SD | 0.980118 |  |  |  |  |  | 0.685181 |
|  |  |  | SEM | 0.565871 |  |  |  |  |  | 0.395589 |
|  |  |  |  |  |  | P value | 0.002577006 |  |  |  |

**Table T6.**
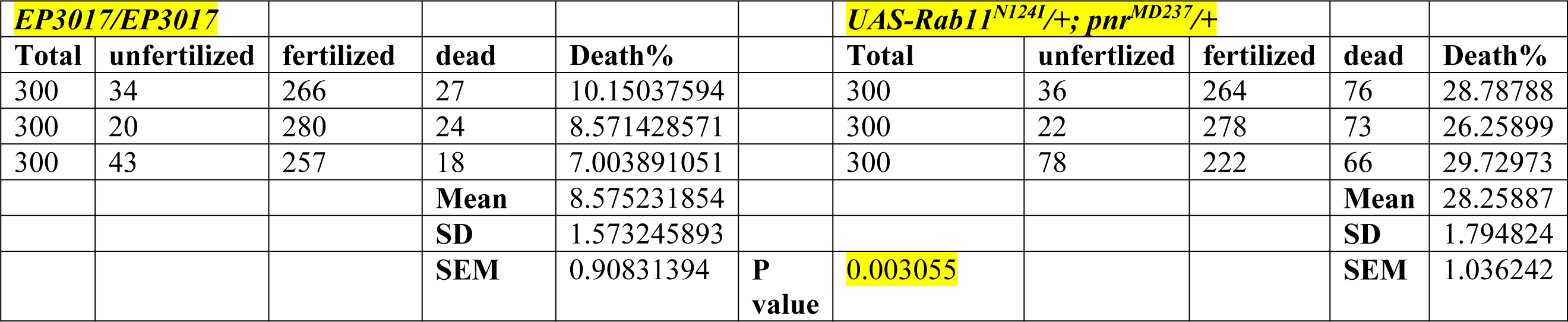
(In support of Supplementary Fig. S1):

**Table T7.**
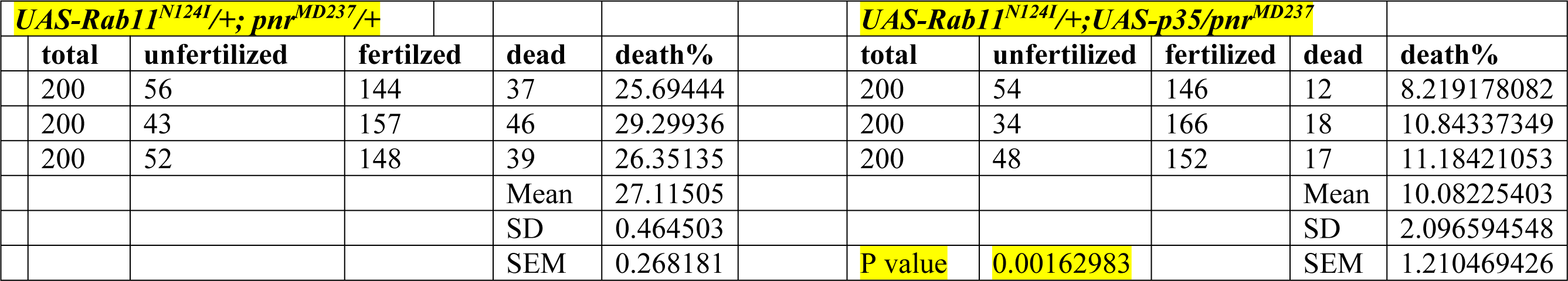
(In support of Supplementary Fig S5)

